# JAK2 signalling to HNRNPA1 represses retrotransposon activity in haematopoietic stem cells

**DOI:** 10.1101/2025.07.07.663405

**Authors:** Jeremy W. Deuel, Andrejs Makarovs, Sheidasadat Goosheh, George Winder, Hisham Mohammed, Jason S. Carroll, Michael Spencer Chapman, Chiraag D. Kapadia, Peter Campbell, Jyoti Nangalia, Elisa Laurenti, Constanze Schneider, Thomas Oellerich, Anthony R. Green, Stephen J. Loughran

## Abstract

Retrotransposon expression must be tightly controlled, particularly in long-lived multipotent stem cells, to prevent deleterious consequences including insertional mutations. Several mechanisms are known to repress retrotransposon transcription during development which are generally thought to persist thereafter. However, integration of retrotransposable elements into host genomes has also provided a major source of genetic variation across evolution. This has generated many sequences that have acquired advantageous host functions, but little is known about retrotransposon expression and mobility in adult stem cells or about how these processes might be dynamically regulated. Here we describe the landscape of somatic retrotransposition in haematopoietic stem cells and identify a previously unrecognised pathway which links cytokine signalling, RNA-modulating HNRNP complexes and repression of retrotransposon activity. Activation of JAK2, by thrombopoietin or gain-of-function JAK2 mutations, triggers tyrosine phosphorylation of HNRNPA1 that represses expression of ERVs, LINEs and SINEs and reduces insertional mutagenesis. This pathway coordinates retrotransposon activity with cellular context and state and provides a mechanism for protecting the haematopoietic stem cell genome.

## Introduction

Sequences derived from retrotransposons litter genomic landscapes. They make up more than 30% of human and mouse genomes and include long and short interspersed nuclear elements (LINEs and SINEs), as well as endogenous retroviruses (ERVs)^1,2^. Transcriptional silencing of retrotransposons during development is necessary to prevent aberrant regulation of neighbouring host genes, activation of anti-viral immune responses, and retrotransposition-driven mutagenesis ^3–8^. Multiple mechanisms are involved in their silencing^9^, many of them imposed at the chromatin level early in development^10^ and thought thereafter to be constitutive, although breakdown of this suppression^11^ can be found in several situations including ageing^4,12,13^, malignancy^4,14,15^ and autoimmune diseases^16,17^. RNA and proteins from these escapee retrotransposons promote pathology by several mechanisms including inflammation-induced premature senescence, viral Env protein-mediated cell fusion, and insertional mutagenesis^8,18,19^.

Retrotransposon expression has also been co-opted to benefit the host with examples including their modulation of chromatin accessibility during early embryogenesis^20^, cortical development of the brain^21^, bone mineralisation and repair^22^; haematopoiesis during pregnancy^23^, and innate immune response signalling^24^. Retrotransposon-derived sequences also account for 37% of mouse and 45% of human enhancers^25^. Although retrotransposon repression is often viewed as constitutive and persistent, physiological roles for retrotransposon activity imply the existence of regulatory mechanisms responsive to cellular states and environments. However such pathways are poorly understood.

Little is known about somatic retrotransposition in normal polyclonal tissues, where it is challenging to detect infrequent events in small proportions of cells. It has been described in human brain and colorectal epithelium^26–29^, but there is limited information on its occurrence in other organs, particularly in rare populations of adult stem cells. The accrual of retrotransposon-mediated insertional mutations is likely to be especially dangerous in stem cells given their longevity and ability to self-renew. Moreover retrotransposon integration depends upon cell division, when nuclear membrane breakdown, chromosome unravelling and chromatin opening all facilitate access to DNA ^30–33^. Nothing is known about mechanisms that might mitigate this vulnerability.

Haematopoietic stem cells (HSCs) are extremely rare bone marrow cells capable of regenerating the whole haematopoietic system and, at steady state, able to generate millions of blood cells every second^34^. Most HSCs are in a quiescent state, while some are activated to divide in response to extracellular cues, including cytokines such as thrombopoietin (THPO)^35–37^. Binding of THPO to its receptor MPL activates the tyrosine kinase JAK2^38,39^, resulting in phosphorylation of multiple substrates including MPL, STAT transcription factors and JAK2 itself^40,41^. JAK2 is also found in the nucleus^42,43^ where its role is less clear but includes modulation of gene expression by phosphorylating histone H3^43–46^ and the chromatin-modifying enzyme PRMT5^47^.

Little is known about the contribution of retrotransposons to the mutational landscape of HSCs. Retrotransposition levels in mouse HSCs have not been reported, and in humans a recent study described widespread LINE retrotransposition in colorectal epithelium, but detected in HSCs from one individual only a single LINE insertion which was thought to have arisen prior to HSC specification^26^. The latter report raised the possibility that the HSC genome may be particularly well protected from retrotransposon activity despite being maintained in an open accessible state associated with multilineage priming and multipotency^48^.

Here we describe the landscape of retrotransposon expression and integration in mouse and human HSCs and identify a previously unrecognised pathway that links cytokine signalling to HNRNP function and retrotransposon activity.

## Results

### The contribution of somatic retrotranspositions to the mutational landscape of human and mouse HSCs

To detect rare somatic retrotranspositions amidst the plethora of preexisting repetitive elements littering the genome we developed a highly specific and sensitive method (PEAR-TREE, Paired Ends of Aberrant Retrotransposons in phylogenetic TREEs). This searches whole genome sequencing data of single-cell derived samples for the coexistence of multiple features that characterise retrotransposition, including target site duplication, paired junction fragments and precise breakpoints (see Extended Data Fig. 1a and Supplementary Method 1).

Twelve somatic retrotransposition events (4 LINEs and 8 SINEs) were detected in 3120 human haematopoietic single-cell derived colonies from seven donors aged from 29 to 81 years old (Fig. 1a,b,c, Extended Data Fig. 1b, Supplementary Table 1)^49,50^. In humans, specification to the HSC lineage occurs in cells that have accrued between 8 and 21 single base substitutions (SBSs)^51^. Nine of the new insertions occurred in cells carrying 21 or more SBSs, whereas two SINE and one LINE could have retrotransposed before HSC specification (Fig. 1d). These data demonstrate that retrotransposition of LINEs and SINEs does contribute to the mutational landscape of human HSCs and that most events occur after HSC commitment. The size of the human HSC population remains in the range of 20,000–200,000 after birth^50^, allowing the number of mutations acquired across the population to be estimated, assuming a constant mutation rate across the lifespan after birth.

**Figure 1.**
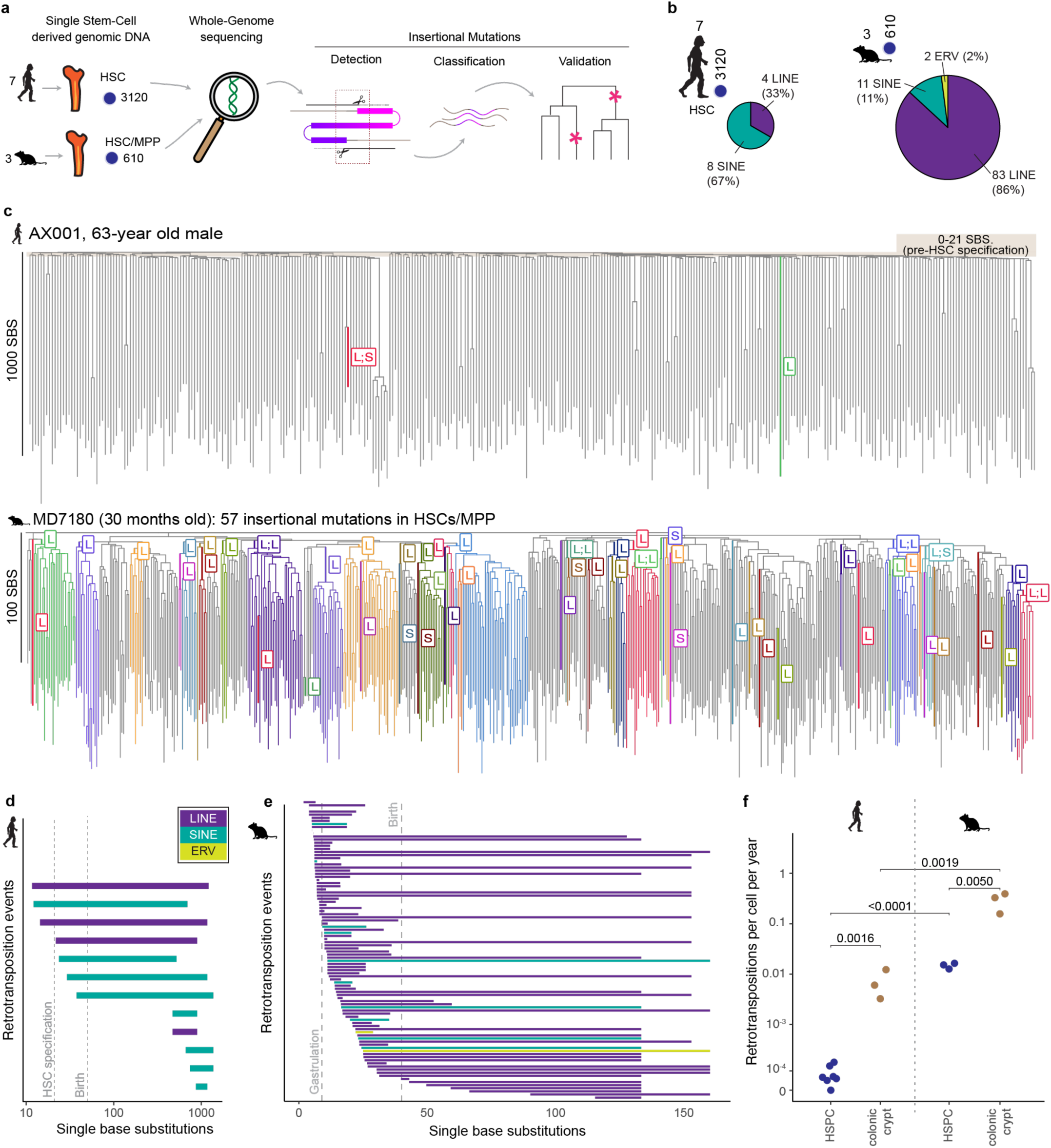
The contribution of somatic retrotranspositions to the mutational landscape of an and mouse HSCs. (a) Detection of somatic mutagenesis by retrotransposons in human and mouse haematopoiesis by PEAR-TREE. Whole-genome sequencing from single stem-cell derived colonies of human haematopoietic stem cells HSCs (7 individuals, 3120 HSCs), and mouse HSCs and multipotent progenitors MPPs (3 individuals, 610 cells) was analysed. Insertional mutagenesis was detected based upon pairs of clipped-reads, classified according to their source retrotransposon and mapped onto HSC phylogenies built from single-base substitutions (SBS). (b) The number of insertional mutations caused by somatic retrotransposition in HSCs of both species (c) Examples of phylogenetic trees inferred from somatic mutations acquired in HSCs of a 63-year old male human donor and a 30-month-old mouse. The tip and edge lengths are scaled to the number of single-base-substitution (SBS) mutations. Each colour represents branches in the phylogeny where a specific somatic retrotransposition was detected, or its presence was inferred from common ancestry. The branch where the mutation was acquired is labelled - L: LINE, S: SINE, E: ERV (d) Plot of 12 somatic insertional mutations detected in seven human donors, indicating the number of SBS mutations that could be present in the cell undergoing each retrotransposition event, inferred by common ancestry in the phylogenetic trees. Colour indicates retrotransposon type. At least nine insertions occurred after the latest possible timepoint of HSC lineage specification (21 SBS, indicated by dotted line), with five of these certainly occurring after birth. (e) Plot of 96 somatic insertional mutations detected in three 30-month-old mice, indicating the number of SBS mutations that could be present in the cell undergoing each retrotransposition event. 5 mutations occurred before the earliest possible timepoint of gastrulation (9-11 SBS; 4 LINEs and 2 SINEs). 85 mutations occurred after gastrulation, including 30 mutations that occurred after birth (40 SBS, 26 LINEs, 3 SINEs and 1 ERV). The timing of 6 mutations could not be resolved relative to gastrulation. (f) Comparison of the rate of insertional mutagenesis caused by retrotransposons in human haematopoietic stem and progenitor cells (HSPC) from 7 donors; human colonic crypts from 3 donors; and from mouse HSPCs and colonic crypts from the same three 30-month-old mice. Colonic crypts had a 94-fold higher insertion rate than HSPCs in humans (95% confidence interval 49-192) and a 21-fold higher insertion rate in mice (95% confidence interval 10.3-40.5). Mouse HSPCs had a 350-fold higher insertion rate than human HSPCs (95% confidence interval 203-763) and mouse colonic crypts had an 80-fold higher insertion rate than human colonic crypts (95% confidence interval 39.3-154). For the calculation of individual rates, see Supplementary Method 2, p-values calculated by *t*-test on logarithmically-transformed data (increasing the minimal number of detected insertions to 1 in one human HSPC sample).

Between birth and the end of the natural lifespan our data predict that humans accrue 460 somatic retrotranspositions within their HSC population (95% confidence interval 126-1456) at a rate of 7.96 × 10^-5^ per HSC-year (95% confidence interval 4.26-13.3 × 10^-5^, see Supplementary Method 2).

Somatic retrotranspositions were more frequent in mice: 96 insertions (83 LINEs, 11 SINEs and 2 ERVs) were detected in 610 single-cell derived colonies from 3 mice aged 30 months^52^ (Fig. 1b,d. Extended Data Fig. 2a, Supplementary Table 2). Each new insertion occurred in a pattern consistent with the phylogenetic tree inferred from shared SBS mutations, confirming the somatic nature of the insertions as well as the precision of PEAR-TREE (99%; Fig 1c, Extended Data Fig. 2a, Supplementary Method 3). The timing of HSC lineage specification relative to SBS burden is not known in mice, but mouse cells undergoing gastrulation have 9-11 SBS, and 6 % of retrotranspositions occurred before this time point. A further 6 % of retrotransposition events could be confirmed to occur in HSCs after birth (40 SBS) while the timing of 88 % of retrotranspositions could not be precisely determined relative to HSC specification. However in humans HSC commitment occurs soon (1-3 SBS) after gastrulation^51^, so it is likely that a large proportion of the murine insertions occur in HSCs (Fig. 1e). Between birth and the 30-month-old time point analysed here, our data predict that each mouse acquires 4610 retrotranspositions in their HSCs (95% confidence interval 1949-10,818) at a rate of 0.0279 per HSC-year (95% confidence interval 0.0193-0.0387; see Supplementary Method 2). Thus murine HSCs accrue 350-fold more insertions per year relative to human HSCs, and 10-fold more across their lifespan.

The increased rate of retrotransposition in mice is much greater than the three-fold higher rate of SBS mutations found in mouse HSCs compared to human HSCs (42-48/year compared to 14-17/year^49,50,53^). The mouse genome contains 20-30 times more functionally-competent LINE elements^54^, and retrotransposition occurs nine times more frequently in the mouse germline (55% of murine compared to 6% of human births)^55,56^. The difference in retrotransposition levels between mouse and human cells is therefore not specific to HSCs, and may reflect, at least in part, the greater burden of active retrotransposons in mice, although our results do not exclude differences in the strength of repressive mechanisms in different host species.

Two of the 12 human insertions arose in the same branch of one phylogenetic tree (Fig. 1c), an event unlikely to be observed if insertional mutations occurred independently from each other (p=0.019, see Supplementary Method 4 for calculation). In mouse HSCs, retrotranspositions also occurred more frequently in HSCs that had already acquired one, leading to accumulation of insertional mutations in certain clades (Fig. 1c; p<0.0001, Supplementary Method 4). This observation could reflect single rogue retrotransposons escaping repression and inserting repeatedly and/or a heritable reduction in repressive mechanisms within certain clades. The latter explanation is favoured by the existence of single phylogenetic branches in which novel insertions were caused by entirely distinct classes of retrotransposon (LINE and SINE), as was the case in the human example and in 2 out of 10 murine phylogenetic branches in which more than one insertion arose (Fig. 1c, Extended data Fig. 2a).

PEAR-TREE analysis of 122 human colonic crypts from three donors (16-56 years old) demonstrated a 94-fold higher rate of somatic retrotransposition compared to HSCs (Fig. 1f, Extended data Fig. 2c). Similarly in mouse colonic crypts, a 21-fold higher rate of retrotransposon events was observed compared to haematopoietic cell derived colonies. As in humans, this increased rate is entirely attributed to LINE elements but not SINEs or ERVs (Fig 1f, Extended data Fig. 2d). The different insertion rates for LINEs and SINEs are surprising, since SINEs require LINE-encoded proteins for retrotransposition and suggests previously unrecognized differences in mechanisms controlling LINE and SINE activity.

Together these results reveal that somatic retrotransposon integrations do contribute to the mutational landscape of mouse and human HSCs with a much higher rate of insertions per HSC-year in mice. HSCs in both species accrue many fewer insertional mutations than colorectal epithelium indicating that HSCs are relatively well-protected from retrotransposition.

### Retrotransposon expression in HSCs is repressed by THPO signalling and by an activating JAK2 mutation

To explore regulation of retrotransposon activity in HSCs, single-cell RNA-seq data was analysed. Substantial levels of ERV, LINE and SINE transcripts were detected in all murine and human HSCs at steady state (Figs 2a-c). Integration of retrotransposons into the genome depends upon host cell division^30–33^ and since THPO receptor stimulation triggers proliferation of HSCs^35–37^ we analysed data from mouse HSCs induced to divide by the administration of THPO^57^. Expression of all three classes of retrotransposon was significantly downregulated in HSCs following THPO treatment (Fig. 2a).

**Figure 2:**
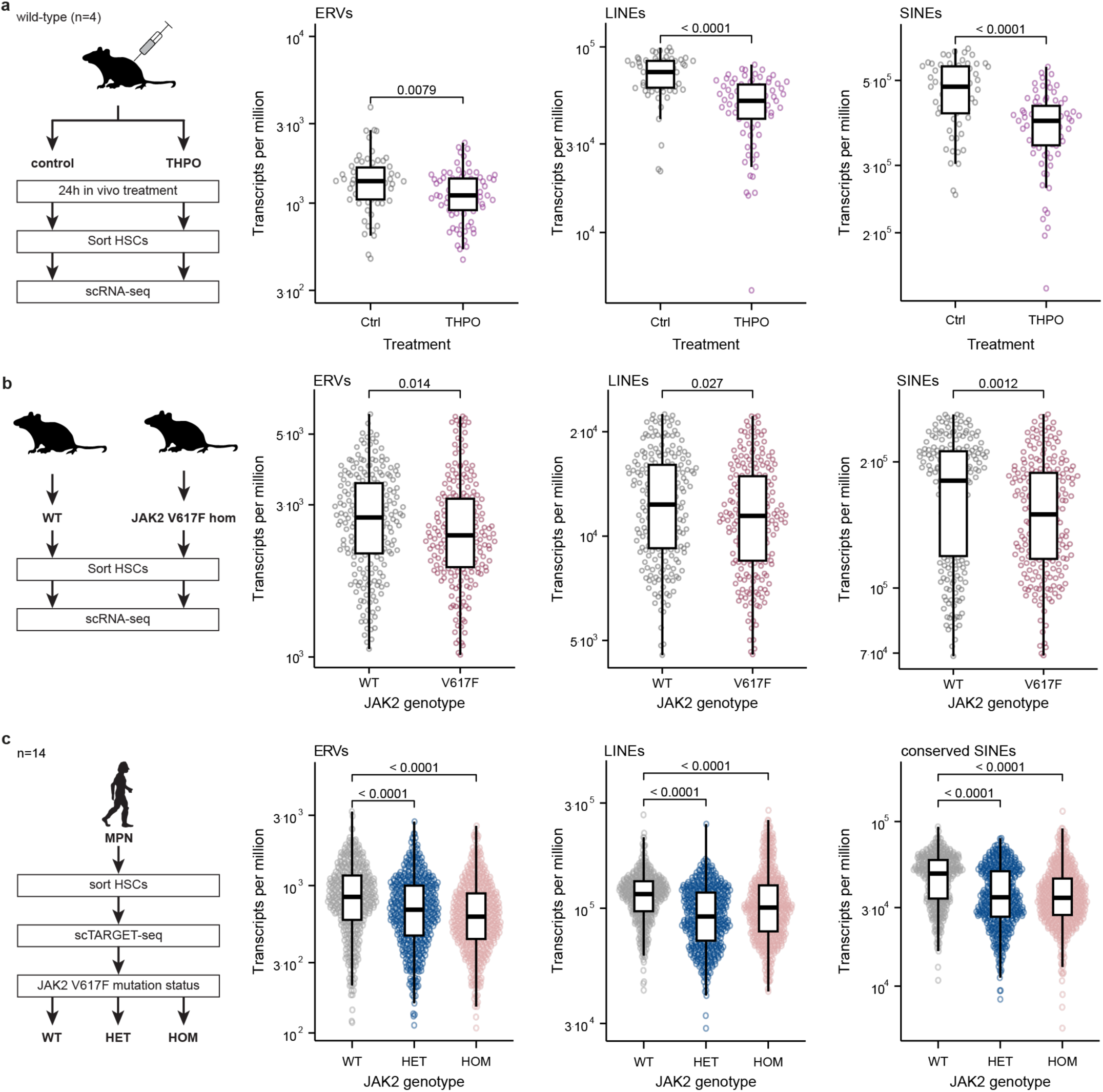
Retrotransposon expression in HSCs is repressed by THPO signalling and by an ating JAK2 mutation. (a) Proportions of retrotransposon transcripts in the transcriptomes of single haematopoietic stem cells (HSCs) after administration of THPO (Nakamura-Ishizu et al.1). Each dot indicates the total proportion of retrotransposon transcripts in a single HSC. RNA-seq was performed 24 hours after wildtype mice were treated with a single dose of THPO or saline (control). ERV=endogenous retrovirus, LINE=long interspersed nuclear element, SINE=short interspersed nuclear element,. (b) Proportions of retrotransposon transcripts in the transcriptomes of single HSCs (ESLAMs, EPCR^+^, Sca-1^+^, c-Kit^+^, CD150^+^, and CD48-) from mice with and without a homozygous activating *JAK2* p.V617F mutation (Kirschner et al2, 192 cells). (c) Proportions of SINEs conserved in human and mice, ERV and LINE transcripts in the transcriptomes of single human HSPCs (Lin-CD34+) from MPN patients (Rodriguez-Meira et al3). *JAK2* p.V617F mutational status of each cell was determined by TARGET-seq (WT, wild-type; HET, heterozygous; HOM, homozygous). All p values determined by Wilcoxon rank-sum test.

Gene set enrichment analysis indicated repression across a broad spectrum of ERV, LINE and SINE elements (Extended Data Fig. 3a). Given the enormous number of retrotransposon loci in the genome, the effects of THPO detected here reflect substantial remodelling of the HSC transcriptome. THPO administration prior to total body irradiation has been shown to protect against radiation-induced LINE activation by upregulating interferon-stimulated genes^58^. However, we found no increased expression of interferon-stimulated genes following THPO treatment (Extended Data Figure 3b), indicating that the retrotransposon repression observed here acts via a different mechanism.

The JAK2 kinase is the key transducer of THPO signalling^38,39^ and we therefore explored the role of JAK2 in retrotransposon repression. ERV, LINE and SINE mRNA levels were all significantly reduced in HSCs from knock-in mice homozygous for the activating JAK2 p.V617F mutation^59^ (Fig 2b). The number of ERV, LINE and SINE loci expressed in each JAK2-mutant cell were also significantly lower, indicating repression acting across a wide range of loci (Extended Data Figure 3c).

Human HSCs from 14 patients harbouring somatic JAK2 p.V617F mutations allowed comparison of JAK2-mutant and wild-type transcriptomes from the same patients^60^. ERV and LINE mRNA was decreased in JAK2-mutant HSCs (Fig. 2c) and the number of retrotransposon loci expressed in JAK2-mutant HSCs was also reduced (Extended Data Fig. 3d). Only a subset of SINEs is conserved between human and mouse; these conserved SINEs were similarly downregulated in JAK2 mutant cells and fewer of these SINE loci were detectably expressed (Fig. 2c. Extended Data Fig. 3d).

These results reveal pervasive ERV, LINE and SINE expression throughout the HSC compartment. Transcript levels of ERVs, LINEs and SINEs in mouse and human HSCs were repressed both by THPO signalling and also by a gain-of-function mutation of JAK2. THPO- and JAK2-mediated repression acted broadly across a large number of diverse retrotransposon loci, resulting in substantial overall changes to the HSC transcriptome.

### JAK2 phosphorylates HNRNPA1

To identify target proteins downstream of JAK2, a phosphoproteomic screen was performed. Three haematopoietic human cell lines with active JAK2 signalling, HEL, L1236 and TMD8^61–63^, were treated with one of two JAK1/2 inhibitors (Ruxolitinib or TG101209) for 30 minutes, and peptides containing phosphotyrosine (pY) were subsequently quantified by mass spectrometry (Fig 3a, Supplementary Tables 3-5). Both inhibitors reduced the amount of multiple phosphotyrosine-peptides by at least one third in each of the three cell lines (Fig 3b). Tyrosine phosphorylation of known JAK2 targets including STAT1, STAT3 and STAT5B were reduced in at least one cell line (Supplementary Tables 3-5). Apart from the JAK2 auto-phosphorylation site JAK2 pY570^64^, only phosphorylation of HNRNPA1 (pY295) was reduced by both inhibitors in all three cell lines (Fig. 3a,b).

**Figure 3:**
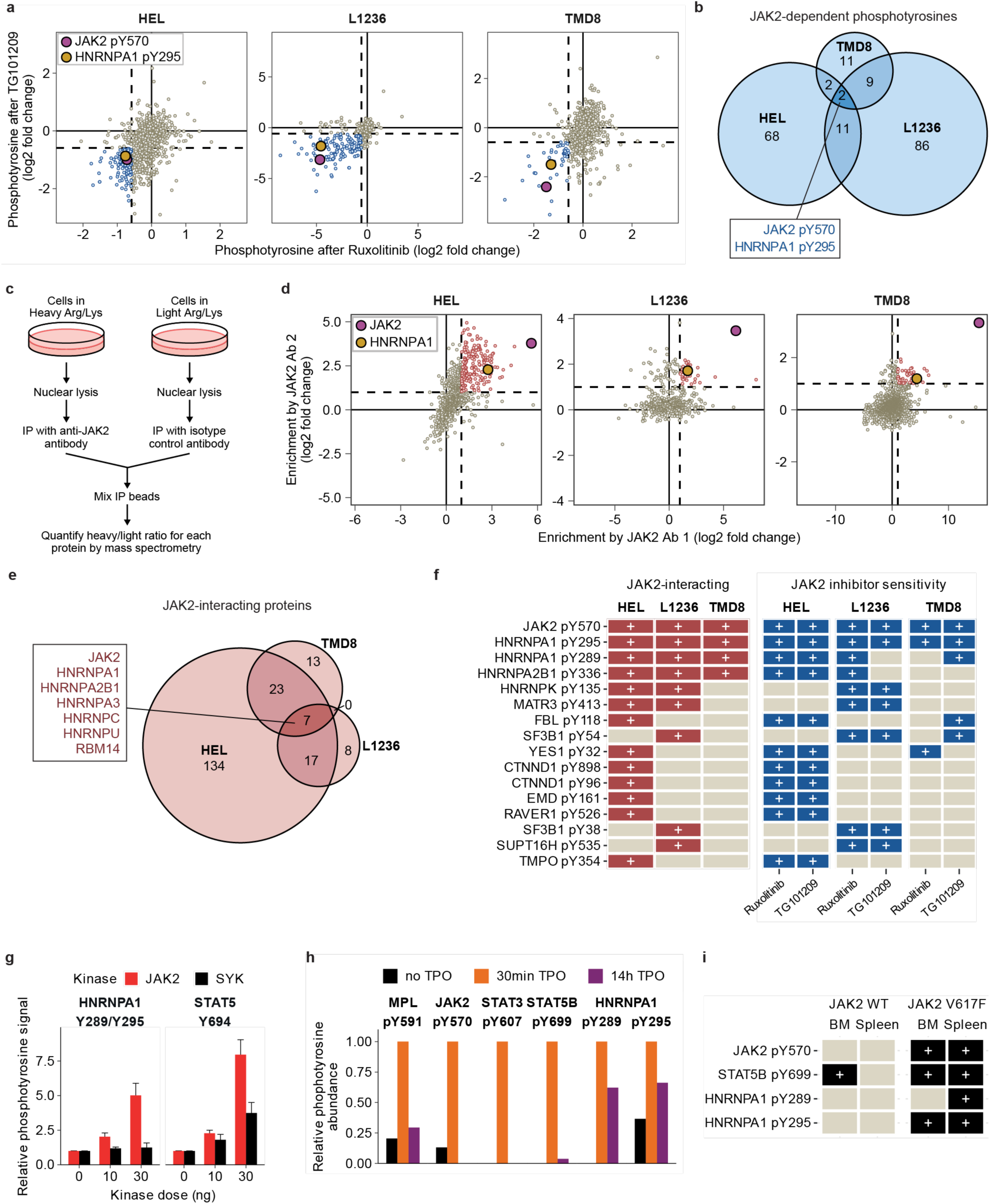
JAK2 phosphorylates HNRNPA1. (a) Phosphoproteomic screening for JAK1/2 dependent phosphotyrosines: Change in phosphorylation levels of tyrosines treated with JAK1/2 inhibitors TG101209 (y-axis) and Ruxolitinib (x axis) of the cell lines HEL, L1236 and TMD8. Phosphotyrosines that were reduced by at least one third by both JAK1/2 inhibitors are shown in blue, HNRNPA1 pY295 and JAK2 pY570 (positive control) are highlighted. (b) Number of phosphotyrosine residues downregulated by at least one third by both JAK1/2 inhibitors in HEL, TMD8 and/or L1236 cells. (c) Experimental setup to screen for nuclear JAK2 interacting proteins, using metabolic labelling. (d) Enrichment of JAK2 interacting proteins in HEL, TMD8 and L1236 cells. Two RIME coimmunoprecipitation experiments are shown for each cell line, using two different anti-JAK2 antibodies. Proteins enriched at least two-fold (dotted lines) by both anti-JAK2 antibodies are indicated in red. JAK2 (positive control) and HNRNPA1 are highlighted. (e) Number of nuclear proteins interacting with JAK2 in HEL, TMD8 and L1236 cells. The seven proteins enriched at least 2-fold by both anti-JAK2 antibodies in all three cell lines are listed. (f) Summary of the JAK2 interactome and JAK1/2 phosphoproteome data. In red, the JAK2 interacting proteins are shown for all three cell lines, (+) signifies enrichment by both JAK2 targeting antibodies. In blue, JAK1/2 inhibitor sensitivity of the phosphotyrosines are shown for both inhibitors and all three cell lines, (+) signifies JAK1/2-dependence of the phosphotyrosines. All phosphotyrosines downregulated by both inhibitors in the same cell line in which the protein interacts with JAK2 are shown. (g) *In vitro* tyrosine phosphorylation of HNRNPA1 and STAT5 peptides by purified recombinant JAK2 and SYK tyrosine kinases, measured by ELISA. (h) Relative levels of phosphotyrosine residues following THPO treatment of 293T MPL cells, measured by mass spectrometry and normalised to levels 30min post THPO exposure. (i) Detection of selected phosphotyrosine residues in primary mouse tissues by mass spectrometry, indicated by a + sign. Tissues from mice homozygous for an activating JAK2 p.V617F mutation, or age- and sex-matched wildtype control mice.

HNRNPA1 is a member of the heterogenous nuclear ribonucleoprotein (hnRNP) family and a core component of multi-protein complexes that associate with nascent RNA polymerase II transcripts to regulate major steps of their processing, including transcription, splicing, stability and transport^65,66^. Several post-translational modifications affect HNRNPA1 function, but regulation by tyrosine phosphorylation has not been previously reported^67^.

Proteins that interact with JAK2 were identified using immunoprecipitation of nuclear extracts and quantitative mass spectrometry as previously described^68^ (Fig. 3c, Supplementary Tables 6-8). Seven proteins were enriched at least 2-fold from all three cell lines by two different anti-JAK2 antibodies: JAK2 itself, and six hnRNP proteins, including HNRNPA1 (Fig. 3d,e).

By combining results of the two screens we identified 16 JAK1/2-dependent phosphotyrosine residues that were also coimmunoprecipitated from nuclear extracts of the same cell line by both anti-JAK2 antibodies (Fig. 3f). These proteins therefore represent candidate direct targets of JAK2. In addition to the known autophosphorylation site JAK2 pY570^64^, only HNRNPA1 pY295 was sensitive to both JAK2 inhibitors and enriched by both anti-JAK2 antibodies in all three cell lines. Levels of HNRNPA1 pY289 were also reduced by at least one JAK inhibitor in all three cell lines (Fig. 3f). Consistent with HNRNPA1 being a direct substrate of JAK2, recombinant JAK2 phosphorylated a synthetic HNRNPA1 peptide in a dose-dependent manner (Fig. 3g).

To explore whether HNRNPA1 Y289 and Y295 are phosphorylated by cytokine signalling, we utilised human HEK 293T cells expressing a THPO receptor (*Mpl*) transgene. Acute THPO stimulation increased phosphorylation of HNRNPA1 Y289 and Y295 together with known JAK2 targets including MPL, JAK2, STAT3 and STAT5B (Fig. 3h, Extended data Fig. 4). In contrast to the canonical JAK2 targets, phosphorylation of HNRNPA1 Y289 and Y295 persisted well above baseline levels after 14 hours of THPO treatment. Although it is challenging to detect tyrosine phosphorylation of JAK2 targets in tissues under physiological conditions^69,70^, we were also able to detect HNRNPA1 pY289 and pY295 in bone marrow and spleen using a mouse line homozygous for an activating JAK2 mutation^71^ (Fig. 3i).

Together these results show that JAK2 phosphorylates HNRNPA1 and that this is increased by THPO stimulation *in vitro* and by an activating JAK2 mutation *in vivo*.

### HNRNPA1 phosphorylation represses retrotransposon expression in HSCs

Tyrosines 289 and 295 lie within the glycine-rich disordered domain of HNRNPA1 (Fig. 4a) and are highly conserved, with Y289 conserved in all vertebrates, and Y295 conserved in all mammals except monotremes (Fig. 4b). To study how HNRNPA1 phosphorylation contributes to the repression of retrotransposons *in vivo*, we generated mice in which tyrosines 289 and 295 were mutated to unphosphorylatable phenylalanine. Three mouse lines were created: a mouse line with a single Y295F mutation (1YF), a mouse line harbouring both Y289F and Y295F mutations (2YF) and a control line that only contained silent mutations necessary for the gene editing (Fig. 4c). All mouse lines were bred to homozygosity, and were viable, fertile and morphologically normal with no differences in litter sizes or sex distribution (Extended Data Fig. 5a,b). No major changes in haematopoiesis were observed in HNRNPA1 1YF and 2YF mice. Haematopoietic stem cell function was normal as assessed by competitive transplantation of bone marrow, and peripheral blood counts were also normal, apart from a modest increase in the fraction of CD8^+^ T cells in older mice (Extended Data Fig. 5c, d).

**Figure 4:**
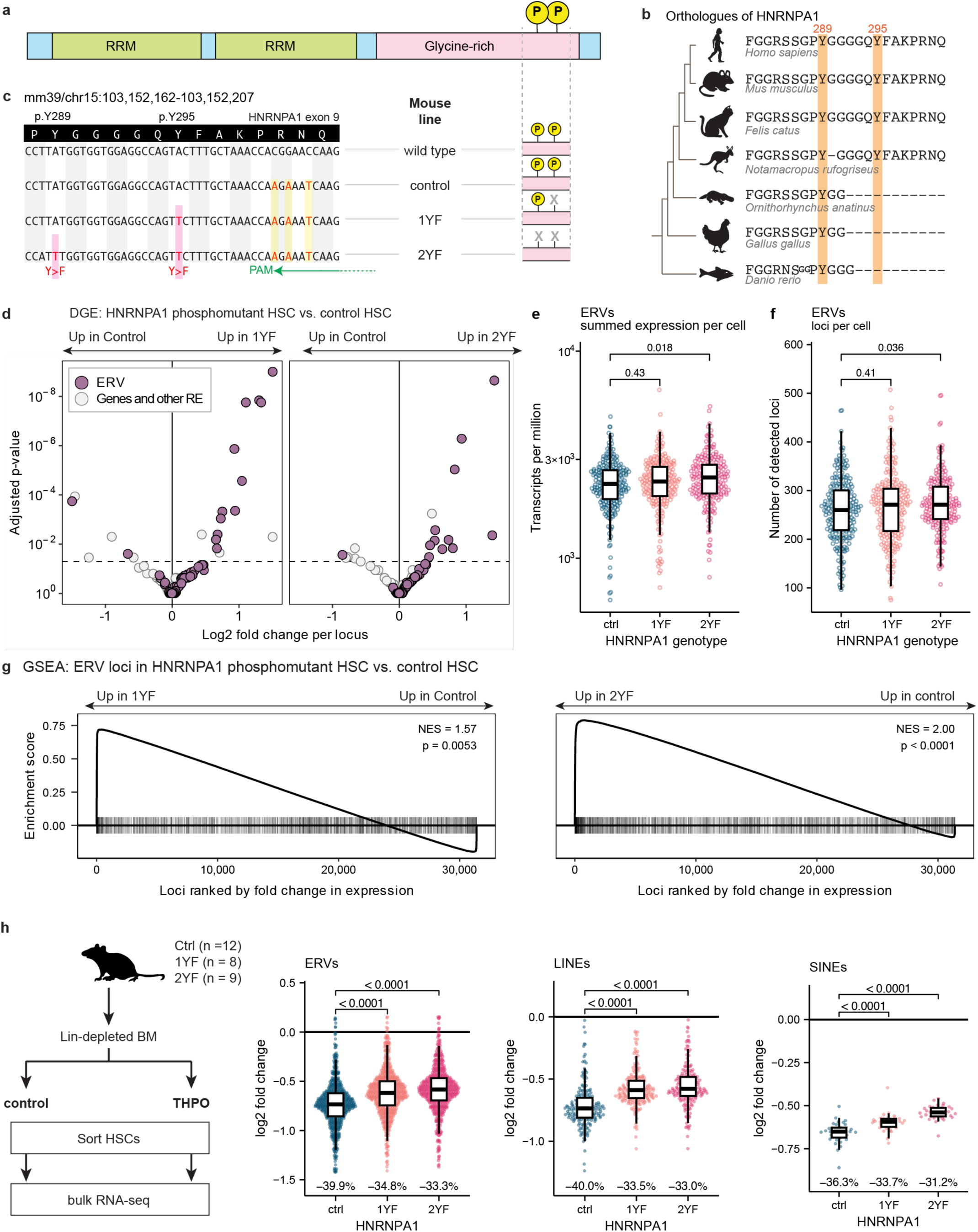
HNRNPA1 phosphorylation represses retrotransposon expression in HSCs. (a) The location of HNRNPA1 pY289 and pY295 are indicated relative to its protein domains. RRM = RNA recognition motif, Glycine-rich disordered domain. (b) Amino acid sequences of human and mouse HNRNPA1 residues 281-302 and homologous sequences from other organisms. Y295 is conserved in the therian subclass of mammals, Y289 is conserved in all examples of the vertebrate subphylum. (c) Three mouse lines were created using CRISPR/Cas9 with a single guide RNA (green): Ctrl which only harboured silent mutations (yellow) and two phosphomutant mouse lines, 1YF with HNRNPA1 Y295F mutation and 2YF with HNRNPA1 Y289F and Y295F mutation (pink). PAM: Protospacer adjacent motif. (d) Volcano plot of differential gene expression (DGE) analysis, of 634 homoeostatic haematopoietic stem cells HSCs (ESLAMS, Lin^−^ CD45^+^ EPCR^+^ CD150^+^ CD48^−^) from 3 adult mice per genotype. Dotted line indicates adjusted p value of 0.05 (Wald test, DESeq2 R package). Purple dots represent endogenous retroviruses ERVs (defined as annotated as “LTR” by dfam.org and flanked by two long terminal repeats) while grey dots show protein-coding genes and non-ERV retrotransposons (RE: retrotransposon elements) (e) The proportion of ERV transcripts in the transcriptomes of each HSC. Transcripts from 17,559 full length ERVs are shown. (f) Number of ERV loci expressed in each HSC. (g) Gene-set enrichment analysis (GSEA) of ERV loci expressed in HNRNPA1 1YF and 2YF HSCs compared to control HSCs. (h) Differential expression analysis of ERV, LINE and SINE transcripts in bulk HSC RNA-sequencing after 16 hours THPO treatment (fold change after THPO treatment is shown). Lineage depleted bone marrow from Ctrl, 1YF and 2YF mice was cultured for 16 hours in the presence or absence of thrombopoietin (THPO). A pool of 100 HSCs (ESLAMs, Lin– CD45+ EPCR+ CD150+ CD48–) was then purified and RNA sequenced. Each dot in the plot represents the fold change of a subfamily of highly sequence-related retrotransposons identified using TEtranscripts^4^ in the THPO treated condition relative to cells cultured without THPO. The median level of retrotransposon mRNA repression is indicated as percentage change at the bottom of the plot. P values of panels e, f and h calculated using Wilcoxon rank-sum test.

We performed single-cell RNA sequencing (scRNA-seq) of highly purified HSCs. This analysis revealed few differences in expression of protein coding genes but identified a significant increase in transcripts from individual ERV loci (flanked by two long terminal repeats) in both HNRNPA1 1YF and 2YF mice (Fig. 4d, Supplementary Tables 9 and 10). The proportion of the transcriptome comprised of ERV mRNA was also modestly but significantly increased in HNRNPA1 2YF HSCs (Fig. 4e), and the number of expressed ERV loci was higher (Fig. 4f) indicating that the increased overall expression in 2YF cells is a broad effect, contributed to by many different ERVs. Gene set enrichment analysis (GSEA) showed significant enrichment of more highly expressed ERV loci in both 1YF and 2YF HSCs (Fig. 4g). We did not find evidence of altered expression of LINEs and SINEs (Extended Data Fig. 6a,b) or of ERV-derived partial fragment sequences in 1YF or 2YF HSCs under steady-state conditions (Extended Data Fig. 6c).

To investigate whether HNRNPA1 phosphorylation is necessary for THPO-induced retrotransposon repression, 1YF, 2YF and control HSCs were analysed by bulk RNA sequencing after overnight culture with THPO or vehicle control (Fig. 4h). Expression levels of ERVs, LINEs and SINEs were all strongly repressed by THPO in control HSCs (regardless of whether multimapping reads were assigned to loci usingTEtranscript^72^ or fractional counts, Fig. 4h, Extended Data Figure 7a). This repression was partially but not completely attenuated in both 1YF and 2YF phosphomutant HSCs, whether compared to control HSCs after culture in the absence of THPO (Fig. 4h) or to uncultured control HSCs (Extended Data Figure 7b,c). Interferon-stimulated genes were not upregulated in HSCs cultured with THPO (Extended Data Figure 7d), which was also the case for HSCs following *in vivo* THPO treatment (Extended Data Figure 3b), further indicating that THPO-mediated repression of retrotransposon transcripts acts via a different pathway. Our results therefore show that tyrosine phosphorylation of HNRNPA1 is required for the full repressive effect of THPO and also indicate the existence of other routes by which THPO represses RTE expression.

Together these data demonstrate that HNRNPA1 phosphorylation is necessary for full repression of retrotransposon expression in HSCs at steady state and also after stimulation by THPO.

### HNRNPA1 phosphorylation represses retrotransposon activity in vitro

To explore a role for HNRNPA1 phosphorylation in repressing retrotransposon-mediated mutagenesis we generated human HEK 293T MPL cells engineered to harbour homozygous tyrosine-to-phenylalanine (YF) mutations of HNRNPA1 Y289 and Y295 (293T/MPL/2YF cells), which preclude phosphorylation of these residues (Fig. 5a). Nine independent clones of edited cells were generated alongside 12 control 293T/MPL clones and these were used in transfection experiments utilising a LINE reporter construct (Fig. 5b)^73^ that results in GFP expression only after retrotransposition (Fig 5c).

**Figure 5:**
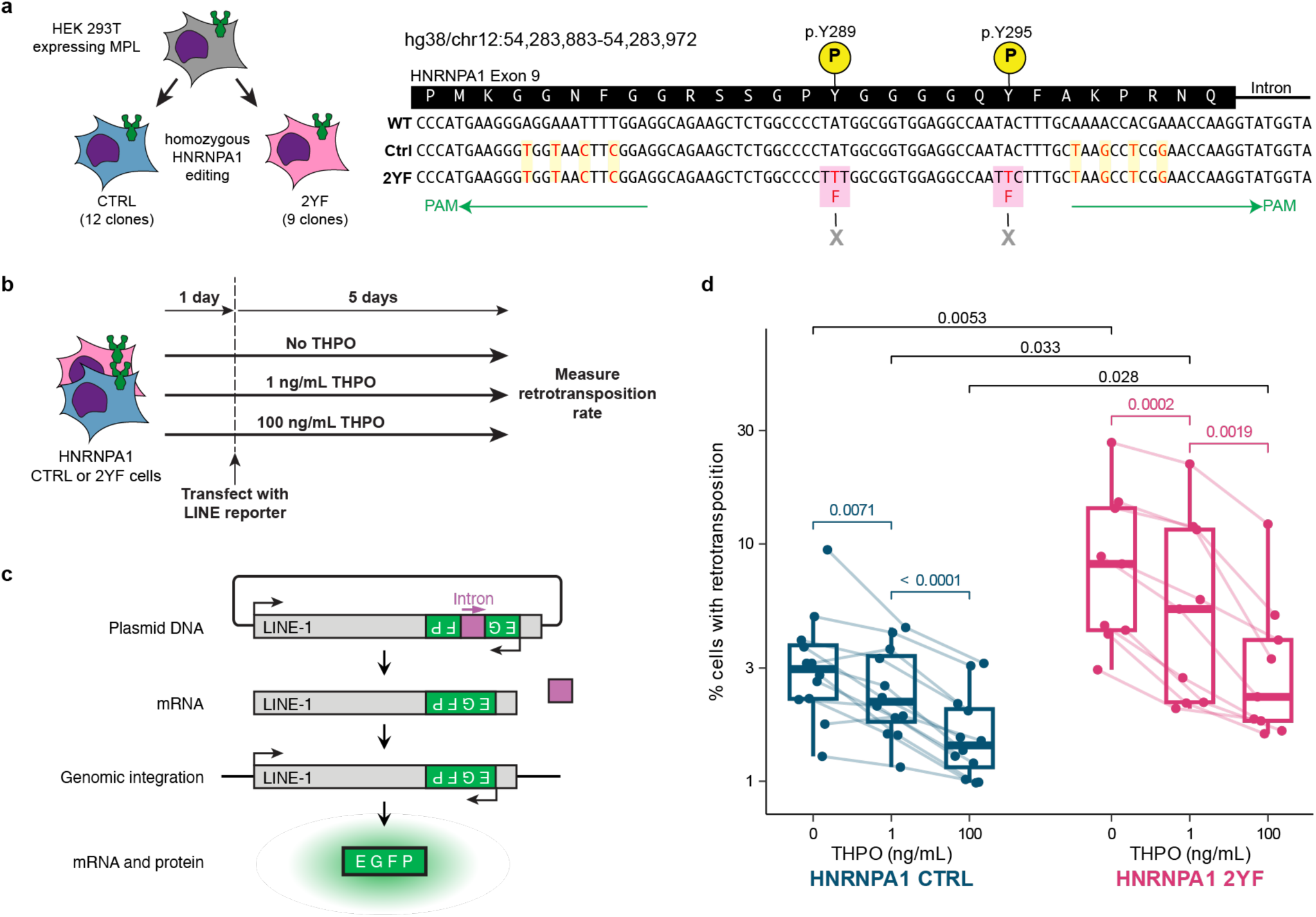
HNRNPA1 phosphorylation represses retrotransposon activity *in vitro*. (a) CRISPR/Cas9 cloning strategy to generate human MPL-expressing HEK293T HNRNPA1 2YF and Ctrl cell lines. The 3’ end of exon 9 of HNRNPA1 is shown, containing the tyrosines (Y) 289 and 295. A double-nickase approach with two guide RNAs targeting the indicated regions (green) was designed. For both Ctrl and 2YF, several silent mutations (yellow highlight) were introduced to prevent re-cutting of already mutated sites. In 2YF, the two tyrosines 289 and 295 are mutated to unphosphorylatable phenylalanine (F) (pink highlight). Both alleles of all cell lines used were homozygously mutated. Nine clonal cell lines of HEK 293T cells harbouring homozygous HNRNPA1 2YF mutations were generated using CRISPR/Cas9 gene editing, alongside 12 clones harbouring only the additional silent mutations necessary for editing. PAM=Protospacer adjacent motif (b) A LINE retrotransposition assay was performed by transfecting HEK 293T MPL cells with a LINE-1 (L1) reporter plasmid and culturing in varying concentrations of THPO. The proportion of transfected cells where LINE retrotransposition had occurred was measured by flow cytometry for GFP. (c) L1 Reporter plasmid EF09R^5^. A LINE1 element (grey) is tagged with a green fluorescent protein EGFP expression cassette (green), encoded on its antisense strand, and interrupted by an intron in the sense orientation (purple). EGFP expression only occurs after transcription and pre-mRNA splicing to remove the intron and subsequent reverse transcription and integration into the host genome to allow transcription of the EGFP ORF from the antisense strand. (d) Percentage of 293T MPL cells with HNRNPA1 Ctrl (silent mutations) and 2YF mutations undergoing retrotranspostion of the L1 reporter in different concentrations of THPO. Each point is the mean percentage from three independent experiments for a 293T MPL clonal cell line generated by gene editing. Lines between the dose levels indicate the percentages of the same clone at different THPO concentrations. P values are shown (Paired *t*-test within (pink/blue) a genotype, unpaired *t*-test (black) between genotypes).

THPO repressed retrotransposition of the reporter construct in a dose-dependent manner in control HEK 293T MPL clones (Fig. 5d). The rate of retrotransposition in 293T/MPL/2YF clones was significantly higher than that of control clones at both doses of THPO (Fig. 5d) demonstrating a substantial contribution of HNRNPA1 phosphorylation to repression of retrotransposition. The THPO dose-dependence of the repression in 293T/MPL/2YF cells suggests the existence of other pathways by which THPO can inhibit retrotransposition independent of HNRNPA1 phosphorylation. The observation that, in the absence of added THPO, 293T/MPL/2YF clones exhibit more retrotransposition than control 293T/MPL clones is likely to reflect either tonic MPL signalling or MPL-independent routes to HNRNPA1 phosphorylation, and is consistent with detectable tyrosine phosphorylation of HNRNPA1 in the absence of added THPO (Fig 3h).

### HNRNPA1 phosphorylation protects the HSC genome from retrotransposition *in vivo*

Next, we investigated whether HNRNPA1 phosphorylation also protects the HSC genome by limiting retrotransposition *in vivo*. Each new somatic insertion is likely to exist in only a tiny subset of cells and so methods (such as PEAR-TREE) that rely on sampling a small proportion of HSC population would not be practical. In designing a new highly sensitive approach we elected to focus on detecting somatic intracisternal A particle (IAP) insertions. This ERV subfamily is most active in the mouse germline and, in contrast to LINEs and SINEs, IAPs transpose their full sequence with high fidelity^54^, facilitating enrichment and precise recognition of sequences at both ends of a somatic IAP insertion. PARTRIDGE (Precise Analysis of Retro-Transposed Replications of IAPs by Deep Genomic Enrichment) uses capture hybridisation to enrich genomic fragments containing the ends of IAPs followed by a sequence analysis pipeline that finds pairs of IAP LTR ends in reads clipped by the aligner (and thus absent from the reference genome). It also identifies the source locus by comparing reads covering the LTR to existing IAPs in the reference genome sequence (Fig. 6a, see Supplementary Methods 5 and 6 for a more detailed description). *In silico* testing demonstrated that PARTRIDGE detects somatic IAP insertions with low allele frequencies with a precision of more than 99% and a recall of 89.6% at allelic frequencies >0.2% (Extended Data Fig. 8a,b).

**Figure 6:**
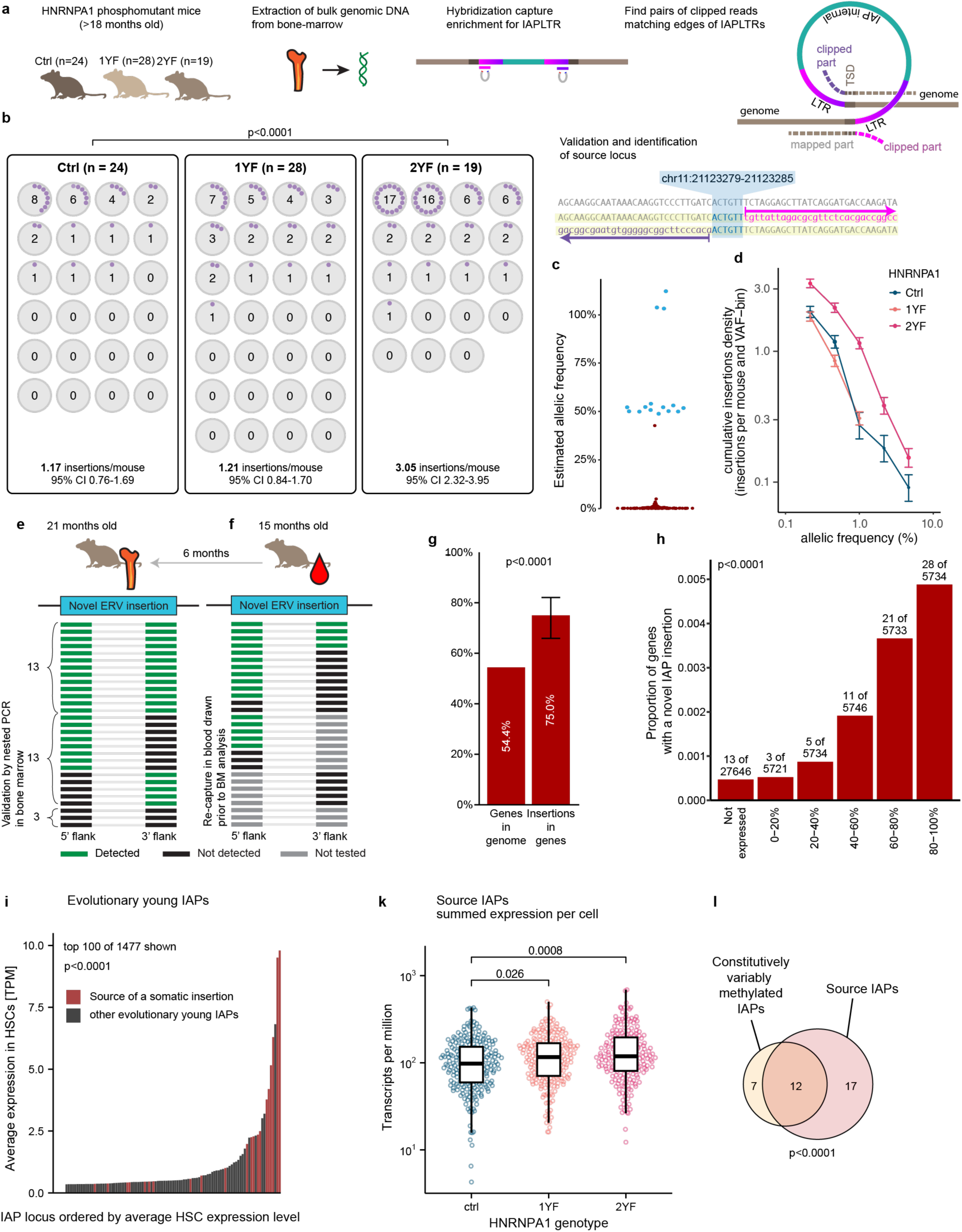
HNRNPA1 phosphorylation protects the HSC genome from retrotransposition *in vivo*. (a) Detection of IAP ERV insertional mutations in bone marrow cells by PARTRIDGE: The 5’ and 3’ ends of IAP LTR genomic DNA was enriched by hybridisation capture and sequenced from the whole bone marrow of seventy-one 21-month-old mice. Insertional mutagenesis was sensitively detected using an algorithm that finds pairs of ends of LTRs within clipped reads within a short window of less than 150bp. The source of novel IAP insertions could often be identified by comparing the sequences of both ends of the inserted LTR to existing IAPs in the reference genome. (b) The number of somatic IAP ERV mutations detected in each mouse. Every somatic IAP insertional mutation (unique to a single mouse) is shown in purple within a mouse (represented by a grey circle). The number of novel insertions per mouse is also indicated. Insertion rates and 95% confidence intervals were determined using Poisson regression. P-Value calculated with Poisson means test (E-test). (c) Estimated allele frequencies (AFs) of IAP ERV insertions detected in a single mouse (red dots), compared to the AFs of a germline ERV insertion (blue dots) detected in multiple mice. The single highest AF of a mutation detected in a single individual was 44.9%, significantly lower than the heterozygous germline insertions with a mean AF of 51.4% and a standard deviation of 1.5%, p=0.002. (d) Cumulative distribution of allelic frequencies of insertional mutations by genotype. Not only is the overall number higher in the 2YF mice, this difference can be seen for every allele frequency (AF) bin, showing higher prevalence also in higher frequency mutations with AF ≥ 1%. The error bars represent the 95% confidence levels of the Poisson model for each AF bin. (e) Validation of somatic IAP ERV insertions in the bone marrow by nested PCR and long-read single molecule Nanopore sequencing. Of the 29 novel somatic insertion sites retested: for 13 (45%), both flanks could be amplified and sequenced, for another 13 (45%) at least one flank could be amplified while for 3 (10%) no PCR product was amplified from either flank. Since 19 out of 58 individual PCR reactions succeeded (33%), 11% of all confirmation reactions are expected to fail for both flanks simultaneously. (f) Long-term persistence of somatic IAP ERV mutations in haematopoietic tissue. Nested PCR and long-read sequencing was performed on DNA from blood drawn from the same mice more than 6 months before harvesting bone marrow. 21 somatic IAP insertions were examined, 17 (81%) were detected. (g) Analysis of 120 destination sites show a preferential insertion into genes. 54.4% of the mm39 mouse reference genome is annotated as a gene in refseq version M36, and 90 of 120 (75%) of novel insertions are within these gene regions, which is significantly more than expected from a random insertion pattern. The confidence interval of the proportion of novel insertions within refseq genes is indicated by the black error bars, the p value was calculated using an exact binomial test. (h) Genes were ranked by their expression quantile in HSCs. The proportion genes in each quantile that acquired a somatic insertions is shown, revealing a significant trend of insertion into genes with higher expression levels (Cochran-Armitage test for trend). (i) Expression of 1,477 evolutionary young (with less than 2 nucleotides difference between both LTRs) IAPEz-loci in HSCs was calculated, thereof the top 100 loci are shown in the plot, ranked by expression level. IAPEz-loci that were identified as sources for novel insertions are coloured red (21 of 29 are in the top 100 loci), while all other analysed loci are shown in grey. Expression of source loci was significantly higher than of other loci. (k) Proportion of mRNA transcripts in single HSC transcriptomes derived from the 29 IAPEz-loci identified as the source of somatic mutations. A significantly higher expression level in HSCs in 1YF and 2YF mutant HSCs was observed. (l) IAPs with constitutive variable methylation (cVM) are enriched in the source IAPs responsible for somatic retrotranspositions. Venn diagram of all uniquely identified ERVs that were the source of somatic mutations (29) and full-length IAPLTR1/2 ERVs loci identified as cVM (n=19) by ^6^ are shown.

PARTRIDGE was used to interrogate bone marrow DNA from a cohort of 71 HNRNPA1 1YF, 2YF and control mice aged 21 months (Fig. 6b, Supplementary Table 11). A total of 120 new insertions were identified with each insertion found in only a single mouse and some mice harbouring multiple distinct insertions. HNRNPA1 control and 1YF mouse lines had similar rates of IAP insertions (approximately one per mouse), consistent with the rate of somatic IAP mutations detected in single-cell derived mouse colonies (Extended Data Fig 8c). In contrast, 2YF mice had a 2.6-fold higher rate of IAP insertional mutations (p < 0.0001). These results show that preventing HNRNPA1 phosphorylation permits an increased level of novel IAP insertions and that precluding phosphorylation of HNRNPA1 Y289, as occurs in the 2YF mice, is necessary for this effect.

All but one of the novel insertions found in single mice had allelic frequencies (AFs) of less than 5% (Fig. 6c) and thus must be somatic. One insertion had an AF of 44.9% and therefore could be germline or be present in an expanded clone – consistent with the latter explanation, its AF was significantly lower than that of multiple heterozygous germ line insertions (Fig 6c; Extended Data Fig 8f). HNRNPA1 2YF mice had a larger number of IAP insertions at all levels of allele frequency (Fig 6d), consistent with a higher rate of insertions occurring throughout their lifespan.

To validate the new insertions using an independent method, nested PCR assays were designed to amplify genomic bone marrow DNA spanning both flanks of 29 of the somatic insertions detected by PARTRIDGE. Sequencing these products confirmed the presence of 26 of the 29 insertions (Fig. 6e, Extended Data Fig. 8d,e, Supplementary Table 12). For 3 insertions neither the 5’ nor the 3’ flanking PCR yielded a product and these remain unconfirmed, an unsurprising rate of PCR failure given the challenges of designing primers to amplify such highly repetitive sequences present at low allele frequencies. Seventeen of 21 tested insertion sites were also detected in peripheral blood DNA obtained from the same mice more than 6 months prior to bone marrow collection, confirming that most new IAP insertions persist in a long lived-bone marrow cell population that contributes to the peripheral blood, consistent with them occurring in HSCs (Fig 6f).

The detection of 120 somatic ERV insertions also provided a unique opportunity to explore patterns of ERV retrotransposition in somatic cells under physiological conditions. Every insertion occurred at a different genomic locus, with at least 300 kb between the nearest two sites (Supplementary Table 11). Eight insertions (6.6%) disrupted exons and, as has been reported for retroviral vectors^74^, significantly more insertions (75%) were observed within genes (Fig 6g) than would be predicted by random site selection. In addition, genes harbouring somatic insertions tended to be highly expressed in HSCs (Fig. 6h). The proviral sequence from 83 novel insertion sites could be accurately mapped to the reference genome, revealing 29 source loci, 13 of which retrotransposed more than once. RNA-seq analysis showed that these source loci were the most highly expressed IAPEz-class ERVs in HSCs (Fig 6i) and they were expressed significantly more in HNRNPA1 phosphomutant HSCs (Fig 6k). Moreover, these 29 mobile IAPs contained the majority (12 out of 19) of a subset of IAPs previously reported to exhibit constitutively variable methylation (cVM-IAPs)^75,76^ (Fig 6l) which are an evolutionary young group of loci which stochastically escape epigenetic silencing during development in different individuals^76^.

The number of IAP insertions detected in each mouse varied more than would be expected if insertional mutagenesis events occurred independently from each other, this was especially true for the 2YF genotype (variance; observed 4.23 vs expected 1.17 for Ctrl; 3.21 vs 1.21 for 1YF; 25.6 vs 3.05 for 2YF). One explanation for this observation could be repeated reinsertions of the same IAP following a single derepression event in some mice. However this would predict multiple somatic insertion events mediated by small number of source loci and did not appear to be the case - in the two 2YF mice with the most IAP insertions, the 17 and 16 insertions originate from at least 8 or 5 distinct source loci respectively. Alternatively, differences in the clonal structure of haematopoiesis between individuals may result in higher allelic frequencies of insertions in some mice, resulting in higher probabilities of detection. Indeed we observed clades of HSCs that are hypersusceptible to retrotransposition in HSC phylogenies (Fig. 1c) and differences in the frequency of these clades between individuals would also account for the observed variance.

## Discussion

Repression of retrotransposon activity is essential to protect eukaryotic genomes from insertional mutagenesis, a hazard that is particularly perilous in long-lived stem cells^77^. Here we demonstrate a direct link in HSCs between cytokine signalling, HNRNP function and retrotransposon activity. The MPL/JAK2/NHRNPA1 pathway described here provides a mechanism for integrating retrotransposon activity with cellular context or state and also for coordinating cytokine-induced proliferation with protection against the increased risk of retrotransposition during cell division^32^.

Despite the risks of retrotransposition, complete and permanent silencing of retrotransposons may be undesirable, since they represent a significant source of genetic novelty and have frequently been coopted to serve important physiological functions for their hosts ^78,23^. Our results reveal high levels of LINE, SINE and ERV transcripts throughout the HSC compartment in mice and humans. This may be a consequence of the accessible chromatin state in HSCs that is associated with their multipotency^48^ and/or reflect a physiological role for retrotransposon expression in HSCs. Consistent with the latter concept, increased expression of retrotransposons in HSCs during pregnancy or chronic blood loss was recently shown to increase erythropoiesis via innate immune signalling ^23^.

However, high levels of retrotransposon expression in HSCs comes with an increased risk of insertional mutagenesis that is particularly acute during cell division^32^ when the nuclear membrane has dissolved, chromosomes are unravelled, chromatin structures are relaxed and DNA is accessible. Signalling-responsive repression mediated by the JAK2-HNRNPA1 pathway provides a way of flexibly mitigating this risk without compromising physiological functions of coopted retrotransposon sequences.

Several lines of evidence point to the importance of retrotransposon-mediated insertional mutagenesis in malignancies: organisms which display the highest retrotransposon expression have higher rates of cancer^79^; approximately 1% of human colon cancers are initiated by somatic retrotransposition events^80^; and leukaemias resulting from retroviral insertions were observed during early trials of HSC gene therapy using gamma-retroviral vectors^81^. Our results predict reduced retrotransposition in MPNs, chronic myeloid malignancies driven by gain-of-function mutations that dysregulate JAK2 activity. Consistent with this idea, analysis of 31 cancer types (mean of 93 patients analysed per type, range = 15-334) revealed somatic retrotransposon insertions in a subset of patients of every cancer type except for MPNs, where no insertions were detected in 46 analysed patients^82^.

This study focussed on the mechanisms of retrotransposon repression in HSCs, but we also found that introduction of the THPO receptor was sufficient to make entirely unrelated human embryonic-kidney-derived cells able to phosphorylate HNRNPA1 and repress retrotransposon activity. JAK2 is activated in multiple tissues by an array of cytokines and receptors^41^ and HNRNP proteins are widely expressed^83^. Moreover THPO-driven repression of retrotransposon activity was attenuated but not completely abolished in the absence of HNRNPA1 phosphorylation, indicating the existence of other repressive pathways downstream of THPO. Indeed, we found evidence of tyrosine phosphorylation of other HNRNP complex components by JAK2 (Fig 3f) including several conserved tyrosines in paralogues such as HNRNPA2B1. Pathways analogous to the one we describe here in HSCs are therefore likely to operate in other cell types to protect their genomes and/or allow dynamic regulation of retrotransposon activity across different tissues, conditions and contexts.

## Materials and Methods

2 Datasets used in this study
| Accession | Species | Description | Reference |
| --- | --- | --- | --- |
| EGAD00001007851<br>EGAD00001006162<br>EGAD00001006118<br>EGAD00001004192<br>EGAD00001004193<br>EGAD00001004086 | Homo sapiens | Whole genome sequencing of blood colonies and laser microdissected colonic crypts | Lee-Six et al 2018<sup>49</sup><br>Mitchell et al, 2022<sup>50</sup>,<br>Chapman, 2021<sup>51</sup><br>Lee-Six et al 2019<sup>84</sup> |
| ERP138320<br>ERP152795 | Mus musculus | Whole genome sequencing of mouse blood colonies | Kapadia et al, 2025<sup>52</sup> |
| GSE121001 | Mus musculus | SMARTer scRNAseq library of haematopoietic stem cells (CD41+ CD150+ CD48- LSK) of mice treated with 100µg/kg thrombopoietin iv. 24h prior to analysis and untreated controls. | Nakamura-Ishizu et al, 2018<sup>57</sup> |
| GSE87631 | Mus musculus | Smartseq2 scRNAseq library of haematopoietic stem cells (ESLAM) of JAK2 V617F homozygous mice compared to wt controls of various age groups. | Kirschner et al, 2017<sup>59</sup> |
| GSE105454 | Homo sapiens | TARGET-seq scRNAseq library of human HSPCs from patients with MPN and healthy controls. | Rodriguez-Meira et al, 2019<sup>60</sup> |
| PRJNA1281824 | Mus musculus | Smartseq2 scRNAseq library of untreated haematopoietic stem cells (ESLAM) of mice with homozygous HNRNPA1 2YF, 1YF and control mutations | This study |
| PRJNA12818<br>54 | Mus<br>musculus | Smartseq2 minibulk libraries of haematopoietic stem cells (ESLAM) treated with THPO or serum-starved with homozygous HNRNPA1 2YF/1YF and control mutations | This study |
| PRJNA12819<br>92 | Mus<br>musculus | Genomic sequencing of peripheral blood of mice with homozygous HNRNPA1 2YF/1YF and control mutations targeted for IAPEz-integration sites | This study |

### Datasets used in this study

#### RNA sequencing data analysis

We created a custom RNA-seq pipeline implemented in Snakemake. The STAR^85^ genome reference was built using the primary assembly of the human GRCh38.v44 or mouse GRCm39.M33 genome (downloaded from Gencode) appended with ERCC92 RNA spike-in sequences. A custom gtf file based on the Gencode annotations and DFAM 3.7^86^ was generated as described below, and used to annotate genes and retrotransposons in the genome. Fastq input files were trimmed using Trimmomatic in the PE or SE setting using the “ILLUMINACLIP” function. Reads were aligned to the reference twice using STAR, using an appended genome for novel splice junctions in the second run. Multimapping reads were included in the final BAM-file. FeatureCounts^87^ was used to count reads associated with annotated features. For each multimapping read, a fractional count of 1/n was assigned to the feature, where n is the number of alignments for that read^87^. For GSEA, interferon-regulated genes were extracted from interferome.com^88^ as previously described^23^.

DESeq2^89^ Version 1.40.2 was used for differential gene expression analysis (DGE). Multimapping counts were converted to integers by using the floor function and the normalisation function of DESeq2 was applied, using the ERCC spike-ins for our own datasets. Only loci with a normalised average expression of at least 1 were considered for DGE. Normalized counts were summed for all loci of a subset of repetitive elements and the logarithmically-transformed values compared between genotype/treatment-groups by Wilcoxon rank sum test. To determine the number of detected loci of a subset of repetitive elements, a locus was considered present if at least one uniquely mapping read could be assigned to that locus. For analysis of retrotransposon subfamilies, we used TEtranscripts with the default settings and its provided GTF annotation^72^. Gene set enrichment analysis (GSEA) was performed with the fgsea package Version 1.26.0 using the DESeq2 log2-fold change in expression as the ranking statistic.

### Annotation of ERV loci

Annotations of repetitive elements, identified using RepeatMasker (https://www.repeatmasker.org), were downloaded from the UCSC genome browser. Putative ERV sequences were identified using long terminal repeats (LTRs) separated by more than 300bp, where more than 25% of the separating sequence was annotated as internal ERV sequence. Total ERV length, including the LTRs, was also required to be longer than 1kb. To select evolutionarily young ERVs, we performed pairwise sequence alignment of the LTRs at the ends of each ERV using the Smith-Waterman algorithm, and measured the similarity of the ends using Kimura’s 2-parameter distance. ERVs with LTR lengths of more than 140bp and a distance of less than 2 divided by the LTR length were considered evolutionary young. By this definition, we identified 17,559 ERVs in the GRCm39 mouse genome, 2,795 of which were evolutionary young. We identified 21,894 ERVs in the GRCh38 human genome, only one of which was evolutionary young (a HERV-H locus at chr6:18754144-18759870).

### Validation of somatic IAP mutations

Four PCR reactions were performed to validate each somatic IAP insertion. Two PCRs amplified a product spanning the 5’ end of the insertion and the other two spanned the 3’ end. For each pair of PCRs, the “inner” PCR targeted a region nested inside the “outer” PCR product (Extended Data Fig 8d). All primers were designed with Primer3, prioritising sequences repeated across the genome as few times as possible. Outer PCR products were amplified from 100 ng of peripheral blood or bone marrow genomic DNA using Phusion High-Fidelity PCR Master Mix with GC Buffer (New England Biolabs). Outer PCR products were purified using SPRIselect beads (Beckman Coulter), then used in a second PCR reaction, amplifying using the inner PCR primers. Production of the expected inner PCR product was confirmed by agarose gel electrophoresis and sequenced using Oxford Nanopore Technology. For all reactions, two genomic DNA samples were used: one from a mouse predicted to have acquired the somatic mutation, and from a control mouse from the same colony. In all instances, the control DNA never generated sequence consistent with the somatic mutation.

### Generation of the HNRNPA1 YF mouse line

CRISPR/Cas9 gene editing of mouse zygotes and derivation of mutant mice was performed as previously described^90^, with these minor modifications: Wild-type SpCas9 (IDT) was dissolved to 1.67 g/L (10.4µM) and sygRNA (5’GAUACUAUACCUUGGUUCCGUGG) (Sigma) to 100µM. A ssDNA repair construct (5’CGATGAAGGGAGGAAACTTTGGAGGCAGGAGCTCTGGC<u>CCATTTGG</u>TGGTGGAGGCCAGTTCTTTGCTAAACCACGGAACCAAGGTATAGTATCTATGACAAAAGACTGATAGTTA) (Sigma) was dissolved in water to 100µM. 0.5µl of SpCas9 at 10g/L was mixed with 0.78µL of sygRNA at 100µM and incubated at room temperature for 15min, then 49µL of microinjection buffer (10mM Tris-HCl pH 7.4, 0.25mM Na_2_EDTA) was added, followed by 0.78µL of ssDNA repair construct at 100µM.

### Genotyping of HNRNPA1 phosphomutant mice

Using OneTaq polymerase master mix (New England Biolabs), a 50µl reaction was set up using the following primers: 5’TCAGACCCTAAGCACTATTGGC and 5’CATTTGGGTTACTGGGTTCAATTA. 30 thermal cycles with an annealing temperature of 58°C and elongation time of 90 seconds were run, yielding a 530bp product (chr15:103,243,559-103,244,088). This product was then Sanger-sequenced using the first primer as the sequencing primer. To screen for mutants before mice were bred to homozygosity, 0.5µl of NlaIV (NEB) was added to the unpurified PCR product and incubated for 30min at 37°C, fragments were analysed by agarose gel electrophoresis. This restriction digestion yields 332 and 198bp fragments for all three mutant mice alleles (Ctrl, 1YF, 2YF), while not cutting the wt allele.

### Flow cytometry of peripheral blood chimerism

20µL of peripheral blood of mice collected by phlebotomy from the V. saphena magna. Red blood cells were lysed for 10min at room temperature in hypotonic lysis buffer (155mM NH_3_Cl, 10mM KHCO_3_, 0.1mM Na_2_EDTA). After decanting lysed red blood cells, cells were stained in a total volume of 50 µL with 0.5µL each of 7-AAD (Thermo, 1g/L), FITC-anti-CD45.1, PE-anti-CD3e, BV421-anti CD19, APC-anti CD45.2, PE-Cy7-anti-Gr1, and APC-Cy7-anti CD11b and measured on an Attune Flow Cytometer (Thermo). Data was analysed using FlowJo v10.9.0 (BD)

### Analysis of peripheral blood counts

Peripheral blood was collected by phlebotomy from the saphenous vein. Cells were analysed by an ABI Vet ABC Hematology Analyser providing a 3-part differential. 10µL of blood was lysed in 250µL hypotonic lysis buffer for 15min at room temperature, then stained in a total of 50µL staining buffer (PBS pH 7.4 with 2% FCS and 1mM Na_2_EDTA) containing 0.5µL of each of the following antibodies: BV421-anti CD19, FITC-anti CD4, PE-Cy7-anti CD8, PE-anti Gr1, AF700-anti CD3e, APC anti CD49b.

### Phosphoproteomic analysis

Enrichment of peptides with phosphotyrosine residues was performed using the PTMScan P-Tyr-1000 Kit (Cell Signalling) following the manufacturer’s protocol. Briefly, Cells were lysed in urea lysis buffer (9M urea in HEPES 20mM pH 8.0, 1mM sodium orthovanadate, 2.5mM sodium pyrophosphate, 1mM 2-glycerophosphate) using sonication. Cysteines were reduced in 10mM DTT for 30min at 55°C, then alkylated for 15min with 10mM IAA at room temperature. The solution was diluted to 2M Urea with 20mM HEPES pH 8.0 and then digested with trypsin at 10mg/L for 16h at room temperature. Peptides were purified using a C18 column and lyophilized. After resuspension in 1.4mL “IAP” Buffer, immunoprecipitation was performed at 4°C for 6 hours followed by a cleanup step with C18 columns as wash buffer. Resulting peptides were eluted in 2×50µL Buffer B. Additionally, a 20µL aliquot of the sample was set apart prior to the immunoprecipitation, mixed with 10% 3% TFA and cleaned up with a µC18 ZipTip (Sigma Aldrich) to correct for overall protein abundance. Peptides were analysed on an Orbitrap Fusion Lumos Mass Spectrometer (Thermo). Spectra were analysed and quantified using MaxQuant v 2.0.1.0.

### Immunoprecipitation mass spectrometry of JAK2-interacting proteins

JAK2-interacting proteins were detected using rapid immunoprecipitation mass spectrometry of endogenous proteins (RIME) as previously described^68^. Nuclei were isolated using a Nuclei PURE prep kit (Sigma-Aldrich) and lysed in RIME LB3 buffer. Two anti-JAK2 antibodies were used – JAK2 Ab 1: JAK2 polyclonal antibody A302-178A (Bethyl laboratories); JAK2 Ab 2: Jak2 D2E12 rabbit monoclonal (Cell signalling technologies).

### In vitro kinase assays

The in vitro kinase activity of recombinant SYK and JAK2 (Life technologies) on biotinylated HNRNPA1 and STAT5 peptides (Pierce) was measured as previously described^91^. Kinases and peptides were resuspended in kinase buffer (60mM HEPES pH7.5; 5mM MgCl_2_; 3µM Na_3_VO_4_; 1.25mM DTT, 20 µM ATP). After 15 minutes at 37 °C, the reaction was stopped by addition of 25 mM EDTA. Peptides were immobilized on streptavidin-coated plates and phosphorylation efficiency determined by ELISA using 1µg/mL anti-pTyr (Millipore,Billerica, USA) and 1:1000 HRPO-conjugated anti-mouse IgG antibodies (Pierce, Rockford, USA).

### Phosphoproteomics of mouse bone marrow

Bone marrow and spleen cells were extracted from three mice homozygous for the activating JAK2 p.V617F^71^ mutation, alongside three wild-type littermate control mice, and erythrocytes were lysed for 15min at room temperature in hypotonic lysis buffer. After one wash in PBS, cells were lysed in urea buffer, then enrichment of phosphotyrosine peptides and mass spectrometry were performed as described above.

### HEK293T MPL cell lines

MPL cDNA with a C-terminal double strep tag was cloned into a lentiviral vector downstream from a CMV promoter, and upstream a C-terminal puromycin resistance cassette connected with an IRES element. Cells were transfected using PEI^92^ and, after 60h of tissue culture, selected with puromycin 3mg/L. Cells were stimulated with 100µg/L THPO for 30min, then lysed in RIPA buffer including c0mplete Ultra (Roche) and PhosSTOP (Sigma-Aldrich) and phosphorylation of STAT3 and STAT5 was confirmed by western blot. For the phosphoproteomics experiment, cells were cultured in SILAC medium (DMEM + 10% dialysed FBS containing 0.115mM stable isotope labelled Arginine, 0.275mM stable isotope labelled Lysine, 1.3mM Proline) for 14 days and exposed to THPO 200µg/L for 30min, THPO 200µg/L for 16h, or not exposed to THPO in serum-free medium. After washing with PBS, cells were lysed in urea lysis buffer

### Gene editing of HNRNPA1 in HEK 293T cells

See Figure 5a for a diagram of this editing design. A Cas9:sygRNA ribonucleoprotein was formed by mixing 11.6 pmol D10A-Cas9 with 6.9 pmol sgyRNA (Sigma) with gRNA sequences 5’AAAACCACGAAACCAAGGUA and 5’UCCAAAAUUUCCUCCCUUCA in a total of 0.625µL Neon Buffer R (Invitrogen) and incubated for 15min at room temperature. ssODNs were resuspended at 23.3µM in buffer R; for a 2YF mutation we used the sequence 5’AACAATCAGTCTTCAAATTTTGGACCCATGAAGGGTGGTAACTTCGGTGGAAGAAGCTCTGGCCCGTTCGGCGGTGGAGGCCAATTCTTTGCTAAGCCTCGGAACCAAGGTATGGTATCTATGTAATTTTGGATAATGT and for the WT control mutations the sequence 5’AACAATCAGTCTTCAAATTTTGGACCCATGAAGGGTGGTAACTTCGGTGGAAGAAGCTCTGGCCCGTACGGCGGTGGAGGCCAATACTTTGCTAAGCCTCGGAACCAAGGTATGGTATCTATGTAATTTTGGATAATGT. 144,000 MPL expressing HEK293T cells were suspended in 12µL buffer R and combined with 0.625µL of the RNP mix, and 0.625µL of the diluted ssODN. 10 µL of this final mix was and 12µL electroporated with a single pulse at 1150 V for 20ms using a Neon electroporation system (Invitrogen). Electroporated cells were resuspended in pre-warmed medium containing 1 mg/L puromycin and 1.2 nM Alt-R HDR enhancer V2 (Integrated DNA technologies). After 3 days, cells were trypsinised and plated in 96-well plates using limiting dilution to ensure single-cell derived colonies. After approximatively 2 weeks, the targeted region was amplified for genotyping using primers 5’ GGACCTTAGGCGCTTAGTTGA and 5’ CAGTTAAAGGGCTTTCTTCCCAG. A diagnostic restriction digestion using DdeI was used to screen for clonal cell lines with edits, and line with homozygous for the desired edits confirmed using by sequencing using Oxford Nanopore technology.

### LINE reporter Assays

The plasmid EF06R^73^ (a gift from Eline Luning Prak, Addgene Plasmid #42940) was modified by removing a triplication of puromycin and hygromycin resistance cassettes and introducing mCherry under control of the CMV promoter. The entire plasmid sequence was verified using Oxford Nanopore sequencing. 200,000 MPL expressing HEK293T cells were plated into 6-well plates under puromycin selection (1 mg/L) to retain MPL expression. THPO (100 µg/L) was added the night before the transfection. The modified LINE reporter plasmid was transfected using the TransIT-LT1 reagent according to the manufacturer’s protocol. Cells were kept in the presence (or absence) of THPO throughout the experiment. Cells were detached using Trypsin/EDTA and GFP fluorescence was determined on a Attune (Thermo Fisher) Flow Cytometer. Non transfected cells were used as negative controls. The fraction of GFP positive cells was calculated by dividing the number of GFP positive cells by the number of transfected mCherry positive cells.

List of antibodies used in the methods described in this manuscript.

| Target species | Target antigen | Label | Manufacturer | Catalog Nr. |
| --- | --- | --- | --- | --- |
| Mouse | CD45.1 | FITC | Biolegend | 294039 |
| Mouse | CD3e | PE | Biolegend | 100206 |
| Mouse | CD19 | BV421 | Biolegend | 115538 |
| Mouse | CD45.2 | APC | Biolegend | 109814 |
| Mouse | Gr1 | PE-Cy7 | Biolegend | 108416 |
| Mouse | CD11b | APC-Cy7 | Biolegend | 101226 |
| Mouse | CD4 | FITC | Biolegend | 100405 |
| Mouse | CD8 | PE-Cy7 | Biolegend | 100722 |
| Mouse | Gr1 | PE | Biolegend | 108408 |
| Mouse | CD3e | AF700 | Biolegend | 100216 |
| Mouse | CD49b | APC | Biolegend | 108910 |
| Mouse | c-Mpl | PE | R&D | FAB1016P |
| Human | JAK2 pY1007/8 | - | abcam | ab32101 |
| Human | STAT5 pY694 | - | abcam | ab32364 |
| Human | pSTAT3 |  | abcam | ab76315 |
| Human | STAT3 |  | cell signalling | 12640S |
| Human | STAT5 |  | abcam | ab194898 |
| Human | beta-Tubulin |  | abcam | ab52901 |
| Human | JAK2 | - | Invitrogen | MA5-24774 |
| Human | JAK2 |  | abcam | ab5832 |
| Human | JAK2 |  | abcam | 4791 |

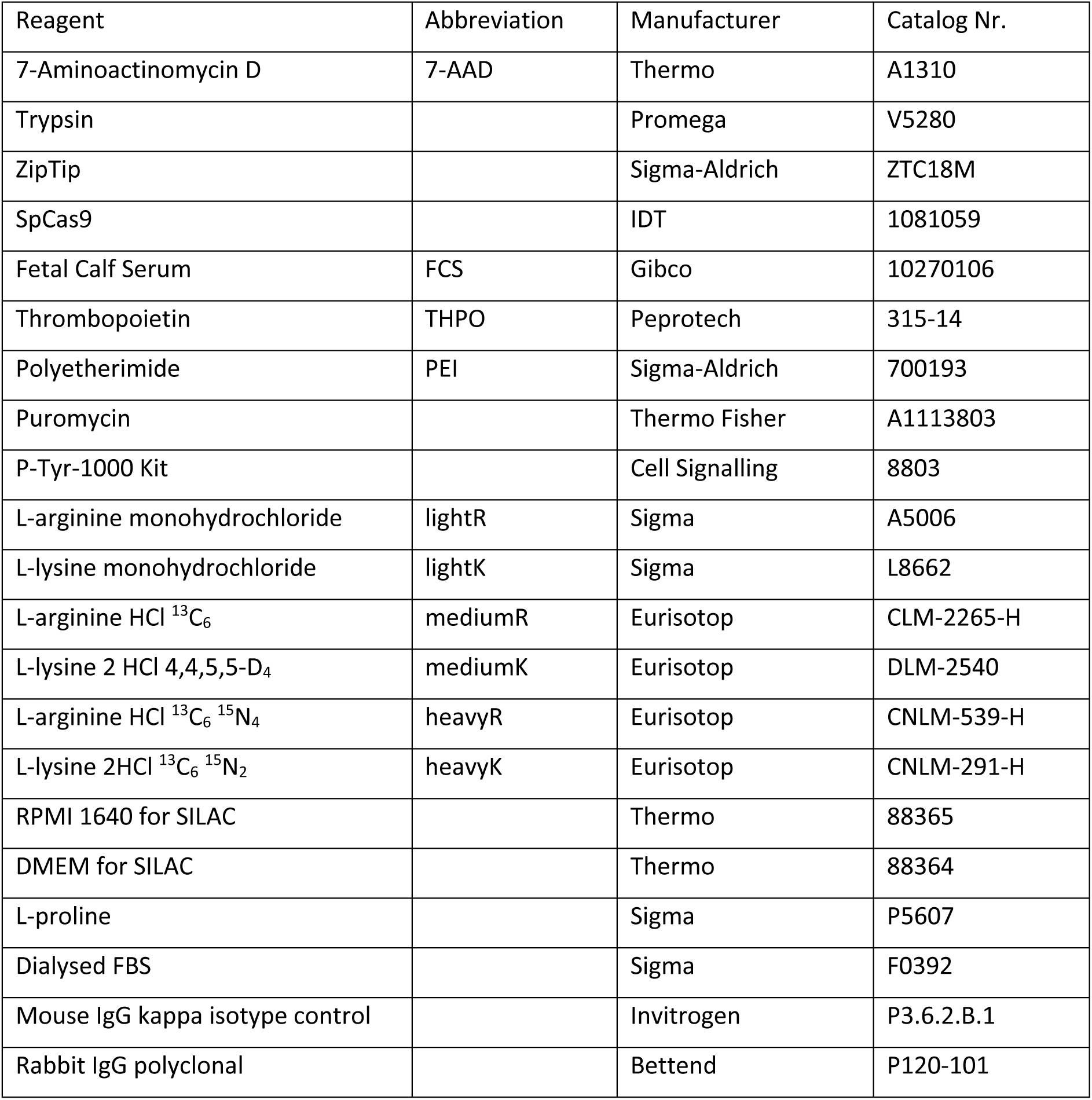
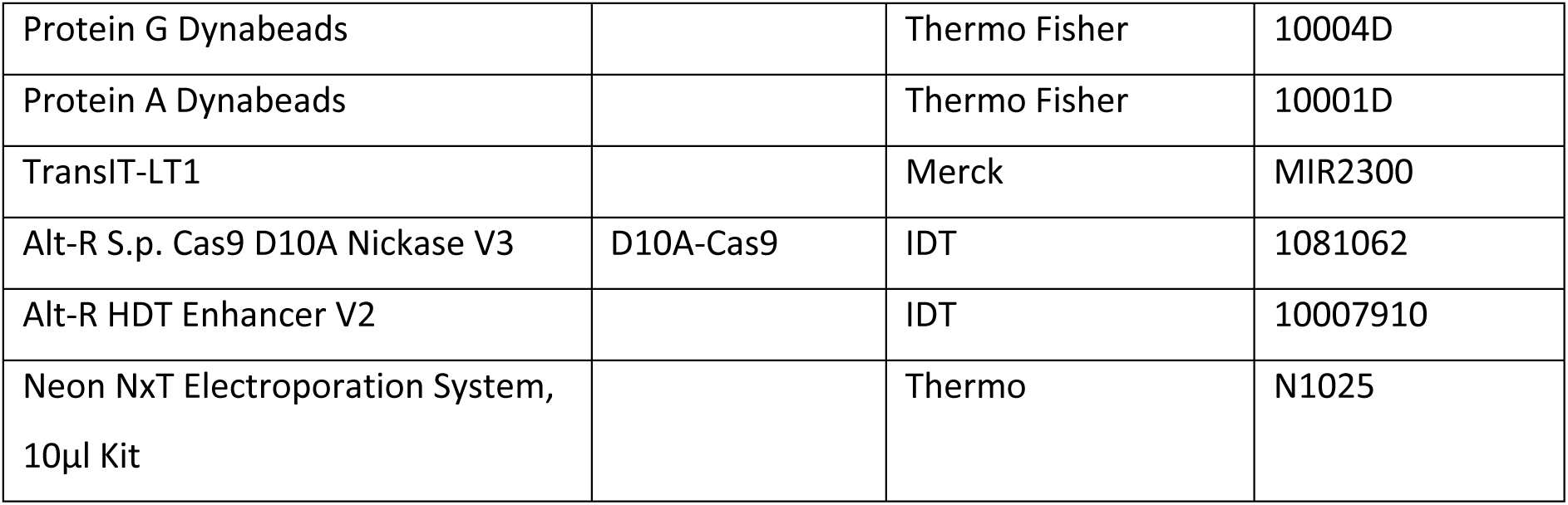
List of reagents and materials.

## Acknowledgements

We thank all technicians in the Green and Oellerich laboratories for their valuable technical assistance: Dean Pask, Tina Hamilton, Juliette Aungier; and Justyna Rak for facilitating approval of mouse experimental work. We also thank Anne Ferguson-Smith, Toshio Suda, Ozgen Deniz, Hugo Bastos, Juan Li and Matthew Williams for their valuable and constructive discussions; This work was supported by Wellcome (RG74909), William B Harrison Foundation (RG91681), Alborada Trust (RG109433), and Cancer Research UK (RG83389) to ARG; as well as the Swiss National Science Foundation (P400PB_183851/1) and the Promedica Foundation (1617/M) to JWD. This research was funded in part by the Wellcome Trust (RG74909). To permit Open Access, the author has applied a CC BY public copyright licence to any Author Accepted Manuscript version arising from this submission.

## Author contributions

JWD and SJL designed and performed experiments and analysed data. AM, SG, GW, HM, JC, CS, and TO performed experiments. JWD developed PEAR-TREE and PARTRIDGE with support from SJL. JWD and SJL performed genetic and phylogenetic analysis with support from MSC, JA, CK, PC and ARG. JWD, SJL and ARG conceptualized the study, interpreted the data and wrote the manuscript. All authors contributed to editing and reviewing the manuscript and approved its publication.

---

**Extended Data Figure 1.**
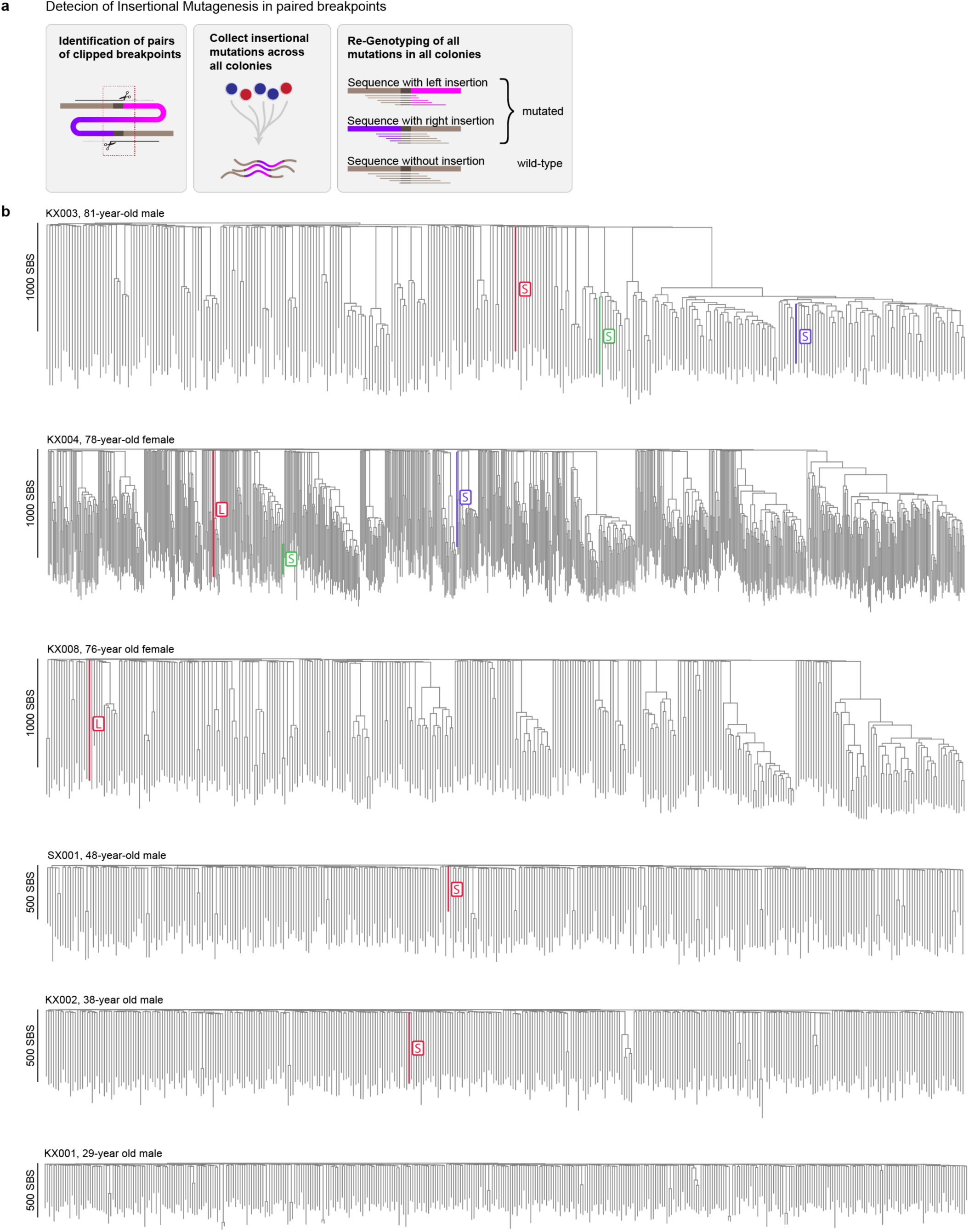
(a) PEAR-TREE - a method to identify somatic insertions: Panel 1: Sequencing reads flanking a novel insertional mutation only partially align to the reference genome (brown), since they do not contain the inserted sequence (pink-purple), thus the non-mapping inserted sequence data is clipped by the aligner. Panel 2: Clipped sequences are grouped by breakpoint within a 40bp window, the break points of the clipped reads must coincide where the alignment to the reference genome ends. Identified pairs of clipped read breakpoints are collected into a single list across all colonies and mice. Panel 3: All colonies are then re-genotyped for all potential insertions to ensure high sensitivity (b) Phylogenetic trees of human HSCs from six donors showing somatic retrotranspositions - L: LINE, S: SINE, SBS: single base substitutions

**Extended Data Figure 2.**
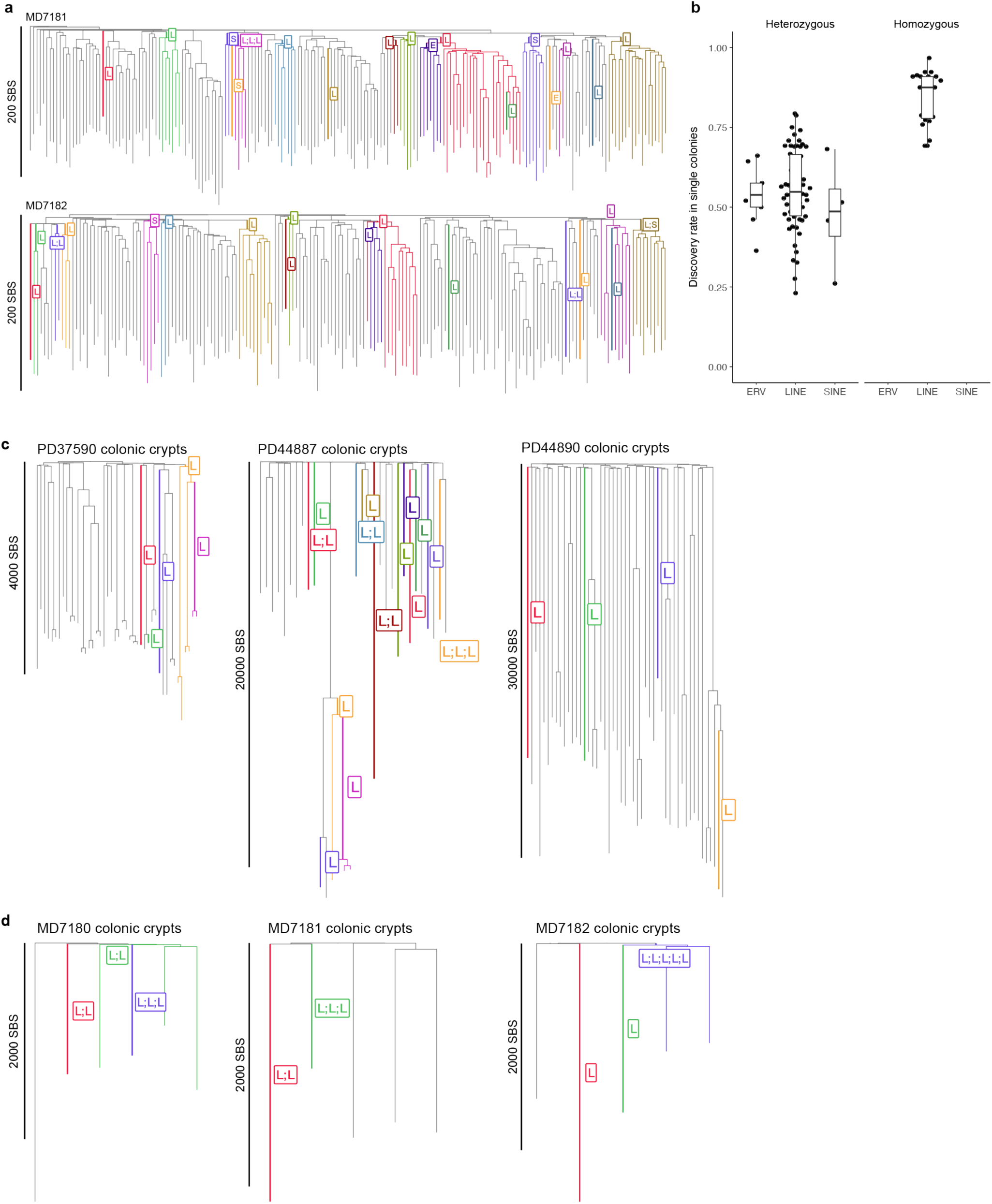
(a) Phylogenetic trees of HSCs from two 30-month-old mice. Each colour represents branches in the phylogeny where a specific somatic retrotransposition was detected, or its presence was inferred from common ancestry. The branch where the mutation was acquired is labelled - L: LINE, S: SINE, E:ERV. (b) Estimating the sensitivity of retrotransposition detection by PEAR-TREE, by determining the probability of detecting heterozygous and homozygous germline insertional mutations in each colony. PEAR-TREE detected 55% of heterozygous insertional mutations in each colony, giving the estimated sensitivity for single-colony insertions (c) Phylogenetic trees of micro-dissected human colonic crypts from three donors with somatic retrotranspositions shown. Each colour represents branches in the phylogeny where a specific somatic retrotransposition was detected, or its presence was inferred from common ancestry. The branch where the mutation was acquired is labelled - L: LINE, S: SINE. (d) Phylogenetic trees of micro-dissected colonic crypts from three 30-month-old mice.

**Extended Data Figure 3.**
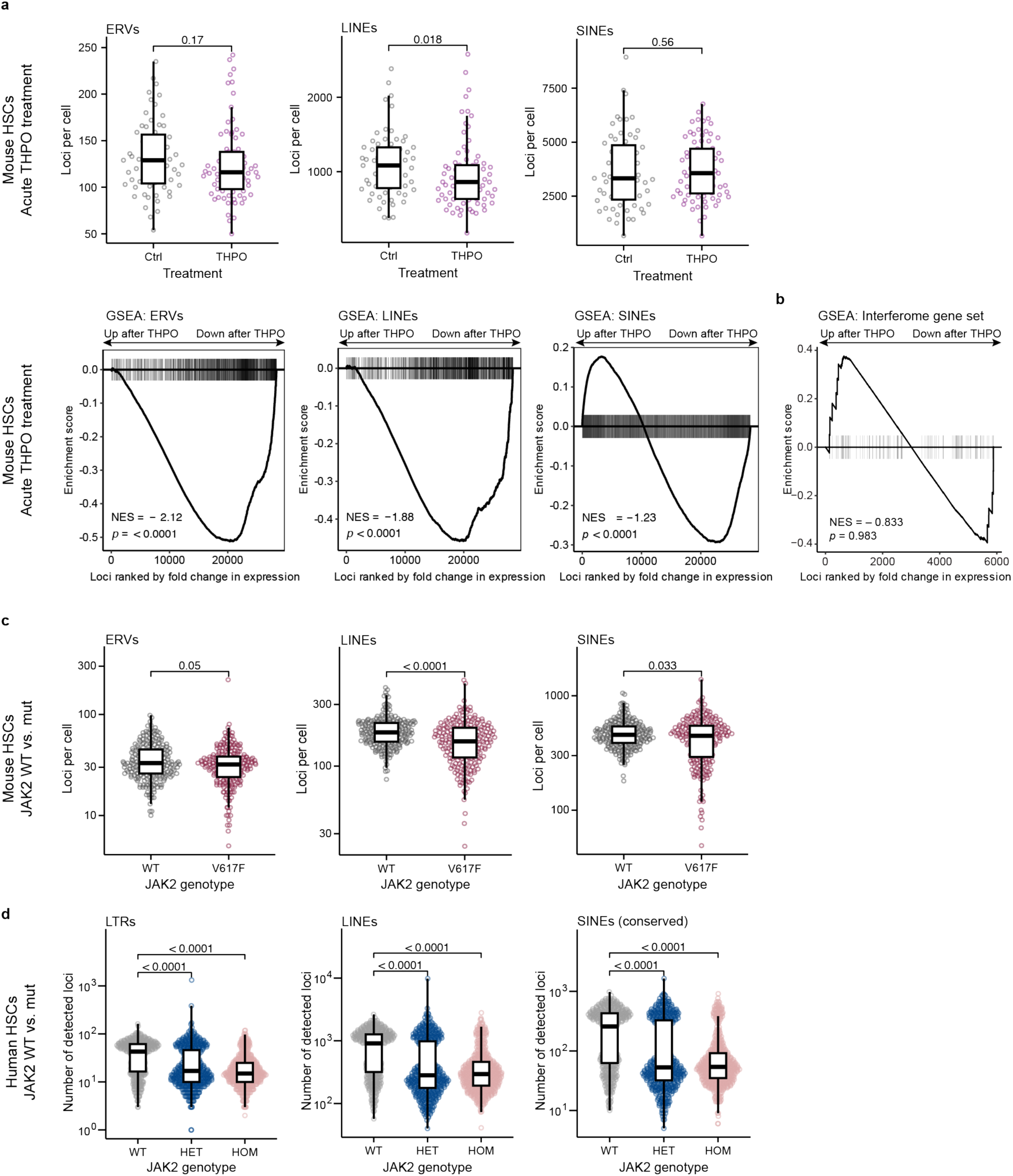
(a) Top row: The number of ERV, LINE, and SINE loci detected in mouse HSC RNA 24h after *in vivo* stimulation with THPO or vehicle control (Ctrl). Each dot represents the number of loci detected in a single HSC. Bottom row: Gene-Set Enrichment Analysis (GSEA) of the three classes of retrotransposons for the same experiment, indicating the downregulation of large numbers of retrotransposon loci. NES=Normalised enrichment score. (b) Gene-Set Enrichment Analysis of interferon-regulated genes after acute THPO treatment showing no significant changes. (c) Number of ERV, LINE, and SINE loci detected in HSC RNA from wild-type (WT) and JAK2 p.V617F homozygous mutant (V617F) mice. Each dot represents the number of loci detected in a single HSC. (d) The number of ERV, LINE and conserved SINE loci detected in RNA from human CD34+ HSPCs. *JAK2* p.V617F mutational status of each cell was determined by TARGET-seq (WT, wild-type; HET, heterozygous; HOM, homozygous). All p values measuring differences in the number of loci detected were determined by Wilcoxon rank-sum test.

**Extended Data Figure 4.**
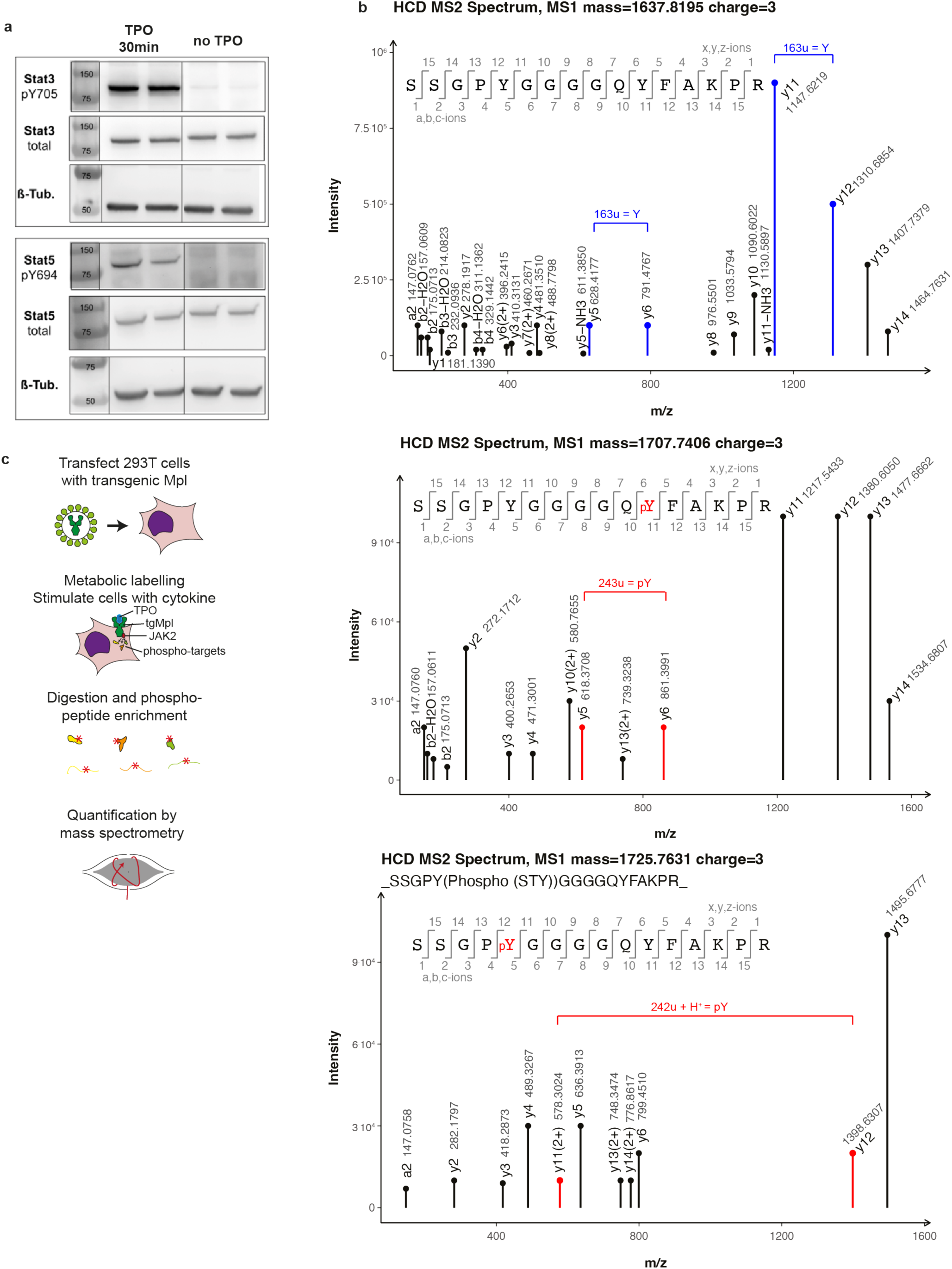
(a) Western-Blots of 293T cells transduced with transgenic *MPL* and stimulated with THPO for 30min. The upper panel shows phosphorylation of STAT3, the lower panel phosphorylation of STAT5. (b) HCD (higher-energy C-trap dissociation) MS2 spectra of HNRNPA1 peptides showing unphosphorylated Y289 and Y295 (top panel), phosphorylated Y295 as evidenced by the y5 and y6 ions, and phosphorylated Y289 as evidenced by the y11(2+) and the y12 ions. The mass difference between two ions of the same charge is expected to equal 163u for a tyrosine (Y) residue and 243u for a phosphotyrosine (pY) residue. The additional charge of the y11 ion is due to the gain of a proton which must be added to the mass difference. (c) Setup for the experiment depicted in Figure 3h. HEK 293T cells were transfected by a lentivirus causing expression of transgenic *MPL* and a Puromycin resistance cassette. After selection, cells were metabolically labelled using stable isotopes of the amino acids arginine and lysine (SILAC). Cells were then treated with recombinant thrombopoietin for 30 min and 14 hours or left unstimulated as controls. After lysis, all cell lysates were pooled, digested and enriched for phosphotyrosine containing peptides, followed by quantification by mass spectrometry.

**Extended Data Figure 5.**
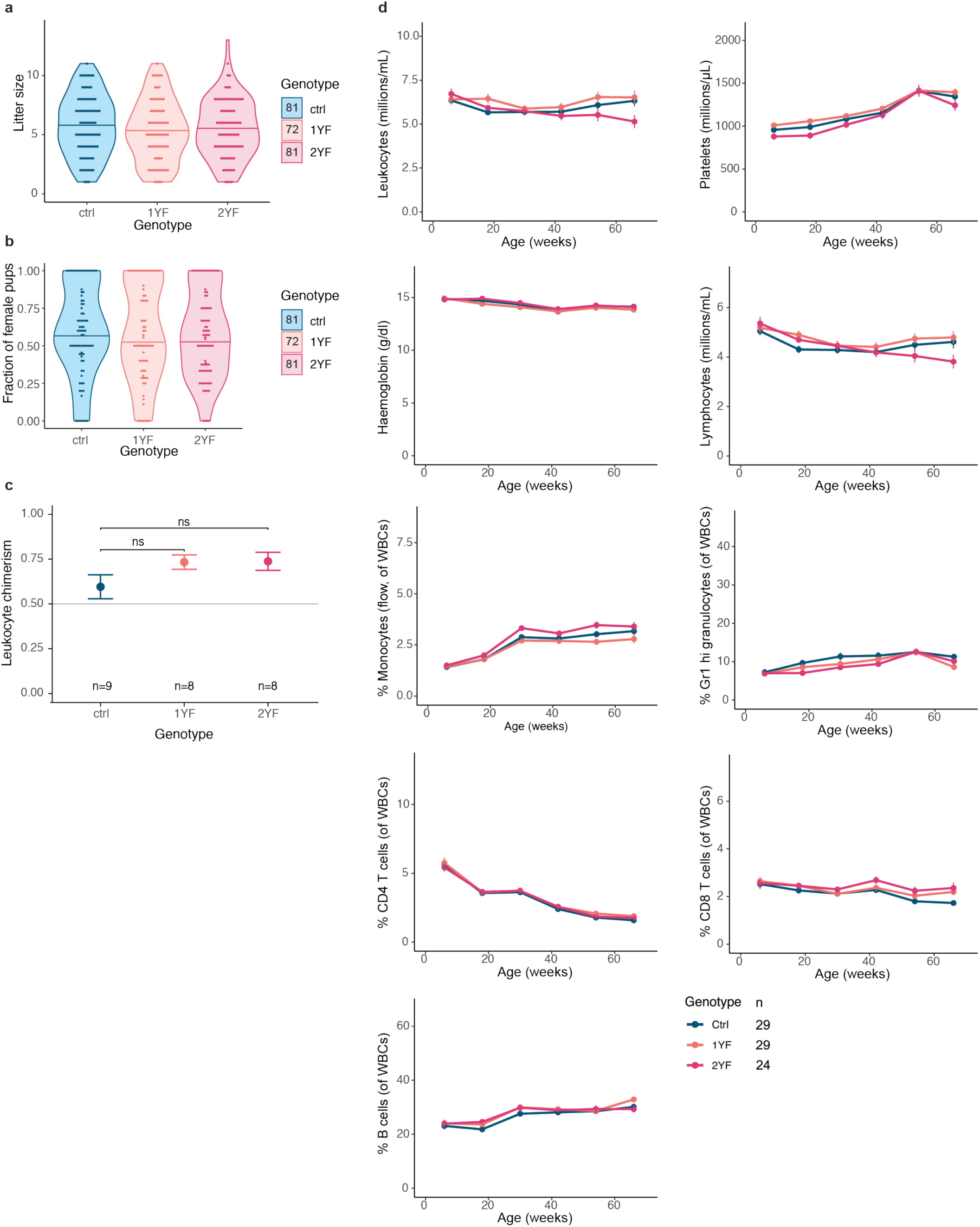
(a) Litter size for 72-81 litters per genotype from HNRNPA1 Ctrl, 1YF and 2YF . No significant differences in litter size were observed. (b) Proportion of male pups in 72-81 litters per genotype. No significant gender bias was observed. (c) Peripheral blood chimerism in recipient mice six months after competitive bone marrow transplantation. (d) Peripheral blood counts and differential counts for all three mouse lines, n=24-29 mice per genotype. Monthly blood counts were summarised in 3-month intervals, the mean and SEM are shown for each data point. No significant differences can be observed between the genotypes except for a higher T-cell count in older 1YF and 2YF mice. P-values in panel b calculated by Wilcoxon rank-sum test.

**Extended Data Figure 6.**
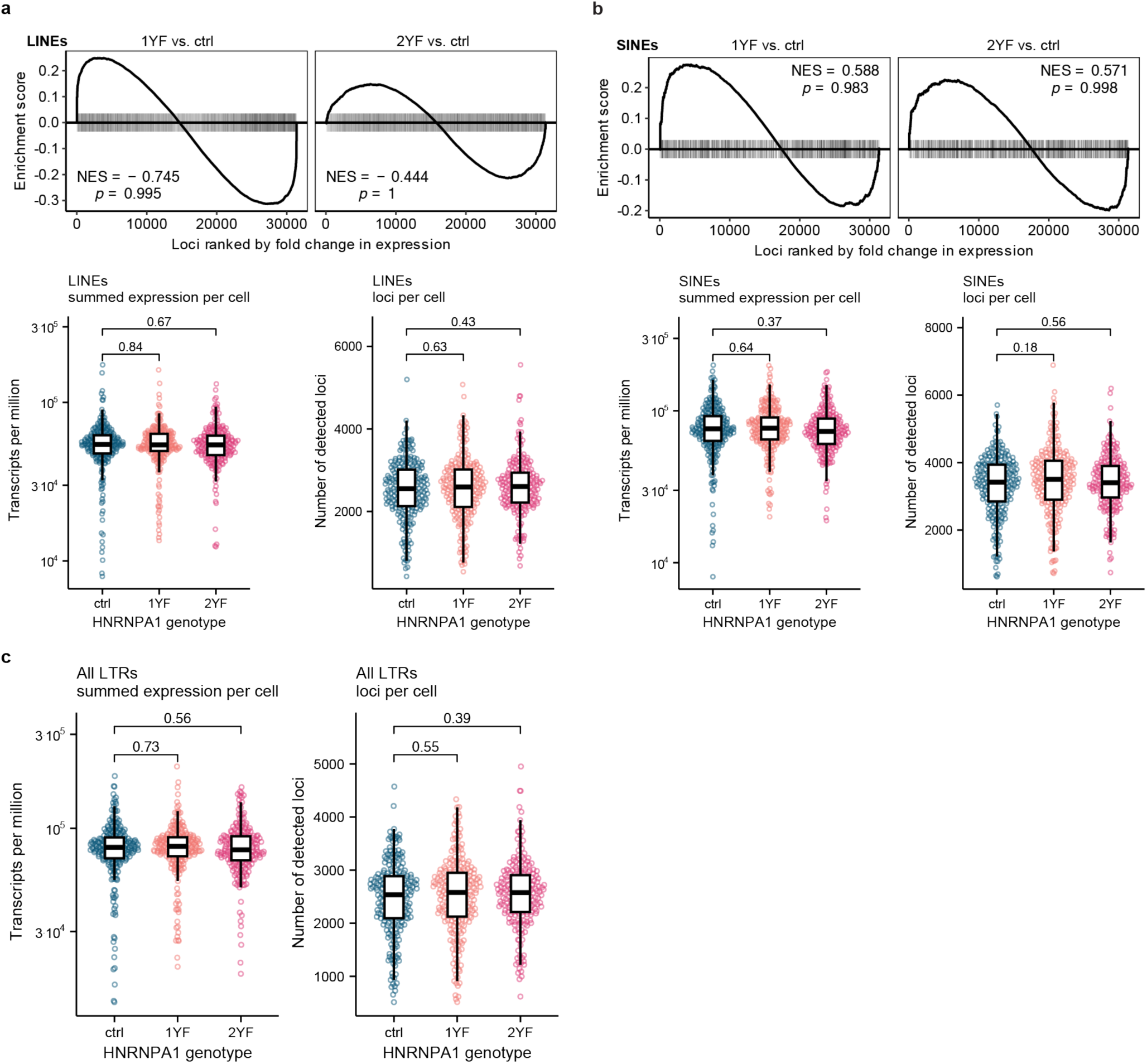
(a) Analysis LINE transcripts in HNRNPA1 phosphomutant HSCs. Top row: Gene set enrichment analysis, bottom row: total expression of LINE transcripts in each HSC (left) and number of LINE loci detected in each HSC (right). GSEA for either 1YF or 2YF when compared to control HSCs did not show any significant enrichment, nor where there any significant changes in total expression or number of detected loci per cell. (b) Analysis of SINE transcripts in HNRNPA1 phosphomutant HSCs. Top row: Gene set enrichment analysis, bottom row: total expression of SINE transcripts in each HSC (left) and number of SINE loci detected in each HSC (right). GSEA for either 1YF or 2YF when compared to control HSCs did not show any significant enrichment, nor where there any significant changes in total expression or number of detected loci per cell. (c) Analysis of all elements classified as “LTR” (long terminal repeat) by dfam.org, including degraded fragments of ERVs and solo-LTRs. There were no significant changes in total summed expression or number of detected loci per cell between the genotypes. p values in panels with dot plots calculated using Wilcoxon rank-sum test.

**Extended Data Figure 7.**
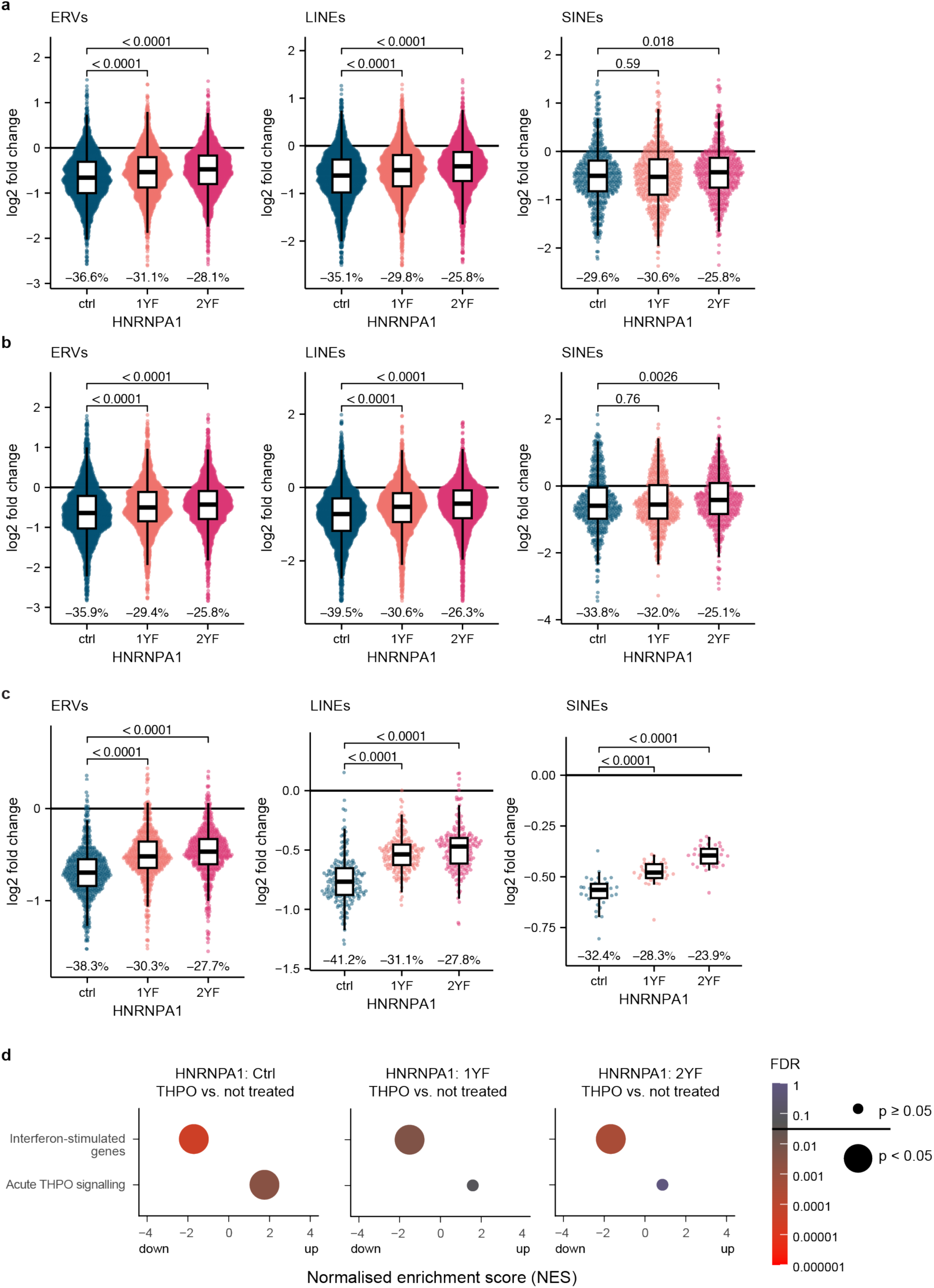
(a, b) Same experiment as shown in Fig. 4h, but using fractional counts to assign multimapping reads to loci instead of using TEtranscripts. In (a), HSCs cultured for 14h in the absence of THPO were used as control, in (b) uncultured HSCs culturing were used as the control condition. Each dot in the plot represents the fold change of an individual retrotransposon locus in the THPO treated condition relative to cells in the control condition. The median percentage change after TPO treatment relative to the control is indicated as a number at the bottom of each boxplot. (c) Same analysis as in Fig. 4h, similarly using TEtranscripts to assign multimapping reads to loci, but using uncultured HSCs as control instead of HSCs cultured for 14h in the absence of THPO. Each dot in the plot represents the fold change of a subfamily of highly sequence-related retrotransposons identified using TEtranscripts^4^ in the THPO treated condition relative to cells lysed prior to culturing. (d) Gene Set Enrichment Analysis (GSEA) of interferon-stimulated genes annotated on interferome.com as well as genes responsive to acute THPO stimulus of cells cultured for 14h with THPO vs uncultured cells. FDR is indicated by colour, gene sets that are changed with an FDR<0.05 are indicated by a large dot. P values of panels a-c calculated using Wilcoxon rank-sum test. The median percentage change relative to the control is indicated as a number at the bottom of each boxplot.

**Extended Data Figure 8.**
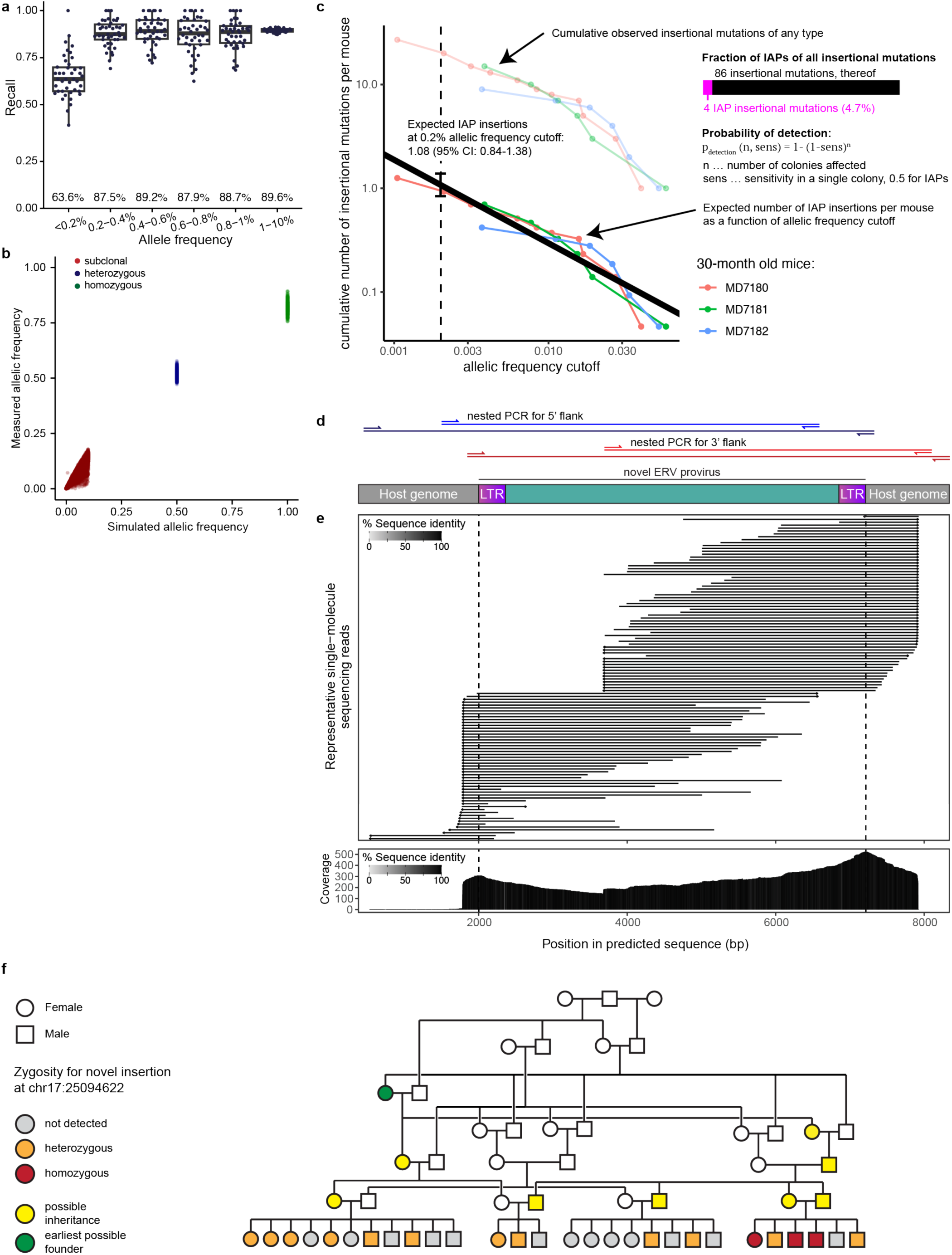
(a) In silico testing of the PARTRIDGE somatic integration detection algorithm. 43 experiments with 10,000 simulated novel insertions each with a randomly chosen simulated allele frequency (AF) of 0-10% were analysed. We were able to detect 89.6% of all simulated novel insertions with >1% AF at a precision of >0.99. Detection levels only start to decrease at AF <0.2%. (b) In silico simulation of the allele frequency of novel insertions calculated by comparing junction read counts mapping to the novel inserted proviral locus, to junction reads counts mapping to the source proviral locus. Heterozygous and homozygous insertions can be well separated, and both have measured allelic frequencies that can be easily distinguished from subclonal insertions (<10% allelic frequency). (c) Estimation of the number of IAP insertions expected to be detected with PARTRIDGE (at a sensitivity of 0.2%), inferred from the PEAR-TREE data. Cumulative allele frequency (AF) of observed insertional mutations were plotted for the three 30-month-old mice (faded coloured lines) and corrected for the sensitivity to detect an IAP as well as the fraction of insertional mutations caused by IAPs (solid coloured lines) An exponential function was fitted, using the number of colonies per mouse as weighting factor. 1.08 insertional mutations per mouse at an AF of 0.2% or higher were expected, 1.17 were observed in Ctrl mice, a difference of less than 10%. (d) Nested PCR validation strategy for novel insertions. The 5’ (blue) and 3’ flank (red) of the novel insertion were targeted individually by nested PCRs. The first PCR (darker colour) had the first primer spanning the insertion point of the opposite flank and the second primer in the reference genome, while the primer pair for the second nested PCR was designed in the reference genome and inside the predicted novel IAP. Inner PCR products were design to overlap from both ends. (e) Alignment of representative sequencing reads from single molecules amplified by PCR. Each horizontal line represents a single amplified DNA molecule matching to the predicted novel ERV sequence. Line colour indicates the sequence identity to the predicted sequence. Additional redundant read alignments that do not span the boundary of the host genome and the novel provirus, or spanning regions already covered by more than 20 other reads have been excluded. In the bottom panel, the total coverage of the predicted sequence of the novel ERV insertion by sequences of single molecules amplified by PCR. Read alignments that do not span the boundary of the host genome and novel provirus were excluded from coverage analysis. The colour indicates the proportion of reads that matched the predicted sequence at each base. (f) Pedigree of the 1YF mouse, showing the inheritance of a germline IAP insertion unique to this mouse colony. The lowest line shows the 28 mice used for IAP insertion analysis, the colour indicates the zygosity for the chr17:25094622-25094628 IAP insertion. A possible inheritance pattern is coloured in yellow; one possible earliest ancestor is coloured in green, showing that this insertion had to occur at least three generations prior to the 28 mice analysed.

