## Supplementary Information for "JAK2 signalling to HNRNPA1 represses retrotransposon activity in haematopoietic stem cells"

### Table of contents

**Supplementary Method 1: PEAR-TREE: Detection of insertional mutagenesis in short-read whole sequencing data** 2

**Supplementary Method 2: Estimation of insertional mutation rates in mouse and human HSCs** 7

**Supplementary Method 3: Calculation of Recall and Precision for PEAR-TREE** 9

**Supplementary Method 4: Testing the independence of insertional mutagenesis events in phylogenetic trees.** 11

**Supplementary Method 5: PARTRIDGE: Detection of IAP-driven insertional mutagenesis in bulk bone marrow DNA** 14

**Supplementary Method 6: Simulation of hybridisation capture enrichment** 18

**References** 19

### Supplementary Method 1:

### PEAR-TREE: Detection of insertional mutagenesis in short-read whole sequencing data

We developed a program, named **PEAR-TREE** (paired ends of aberrant retrotransposons in phylogenetic trees), to detect insertional mutagenesis in short-read whole genome sequencing data. The source code of this program is available on github (https://github.com/jeremydeuel/PEAR-TREE), alongside a description how to install and run the program.

The program was run on samples aligned by bwa-mem2[1] to the human genome (GRCh38) and the mouse genome (GRCm38), in individual bam files (sorted by coordinates) for each colony.

PEAR-TREE works in four successive steps:

1. Discovery of novel insertional mutations,
2. collection and filtering of novel insertional mutations,
3. re-genotyping of filtered novel insertional mutations,
4. collection and final filtering of re-genotyping, and
5. annotation of the novel insertions.

Steps (1) and (3) are applied separately for every bam file, while steps (2), (4), and (5) are applied to all samples from an individual (for human samples) or from a group of related individuals (for mouse samples).

By convention, all alignments in a bam file are relative to the + strand, thus fragments aligning to the - strand are reverse complemented in the bam file. Sequencing reads with an end soft-clipped by the aligner were designated “right clipped” or “left clipped”, depending on which end is clipped relative to the reference genome (Supplementary figure S1).

Right clipped: mapped to the + strand of the reference genome with soft-clipped bases to the right (3' on the + strand or 5' on the - strand), potentially indicating the 5' end of a retrotransposable element if inserted on the + strand or the 3' end if inserted on the - strand.

Left clipped: mapped to the + strand of the reference genome with soft-clipped bases to the left (5' on the + strand or 3’ on the - strand) indicating the 3' end of a retrotransposon on the + strand or the 5' end of a retrotransposon on the – strand.

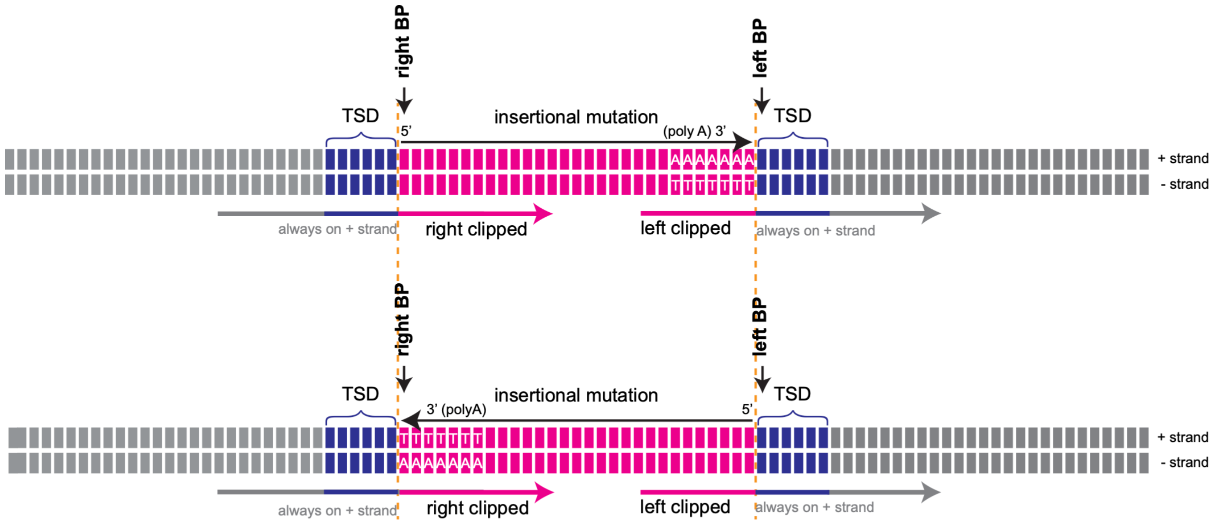


***Supplementary Figure S1: Schema of an insertion in the genome.***

*A novel insertional mutation (most have a polyA tail, as shown in the figure) is depicted in pink. A variable number of flanking bases are duplicated during retrotransposition, called target site duplication (TSD), shown in blue. Read (arrows belows) are mapped to the reference genome (grey) and TSD (part of the reference genome, blue), then clipped either to the right or left. The right breakpoint (BP) is defined as the first clipped base to the right, the left BP is defined as the first mapped base to the left. The top of the figure represents the insertion of a retrotransposon coded on the + strand, the bottom shows an insertion in the opposite orientation.*

The "right breakpoint" (RBP) is the 0-based coordinate of the first non-aligned base of a "right clipped" fragment. The "left breakpoint" (LBP) is the 0-based coordinate of the first aligned base in a "left clipped" fragment. A target site duplication (TSD) can be recognised if coordinate of the RBP is larger than of the LBP, where the length of the TSD = RBP-LBP (See Figure S2). If the coordinate of the LBP is larger than of the RBP, LBP-RBP bases were deleted during insertion, which occasionally has been described for retrotransposon insertions.

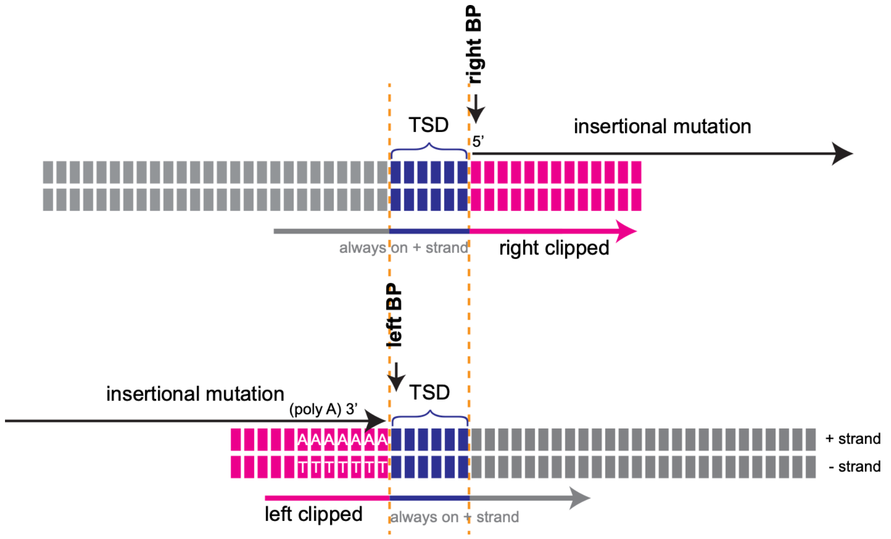


***Supplementary Figure S2**: Situation of the same insertion shown in the top panel of Supplementary Figure S1, as seen by the aligner with respect to the reference genome. The length of the TSD is [right BP] - [left BP].*

**Step 1: Discovery**

Pysam (https://github.com/pysam-developers/pysam) was used to iterate through all fragments present in a bam file. Alignments with a mapping quality (mapq) of less than 60, duplicates, or failing QC checks of the aligner were discarded. Fragments with soft-clipped bases of either end of the fragment, were further considered, others were discarded. Fragments with supplementary alignments (marked as chimeric by the aligner), where any of these supplementary alignments mapped within 1 kb of the main alignment were also discarded, as this is a common artefact produced during sequencing library preparation, most likely caused by the formation of cruciform DNA structures[2]. Next, any part of a soft clipped sequence that resembles adapter sequence or dark sequencing cycles was removed. Only clipped reads with at least 12 bp of remaining soft clipped sequence are further considered. A breakpoint was called if at least two independent fragments cover the insertion point and have identical sequences.

Quality filtering of breakpoints was performed: Breakpoints with homopolymers of more than 6 bases in the aligned part immediately adjacent to the breakpoints were removed, since the determination of the breakpoint was no longer possible. Regions of the genome with more than 120x coverage were removed, since they were prone to artefacts. Alignments to non-canonical contigs and mitochondrial DNA were also excluded.

Pairs of LBPs and RBPs within 40 bases are then formed and saved in a text file format. If multiple pairs are possible within this window, all are discarded. Read mates spanning into the novel insertions are extracted from the bam file and the clipped sequences are extended where possible using these mate reads.

**Step 2: Combination of discovered insertions**

All insertions found during step 1 were combined and duplicates resolved. For each breakpoint, the combined aligned and clipped parts of reads were re-mapped to the T2T genome (hs1 for human samples and mMusMuc1.1 for mouse samples) and discarded if any match was found, since these were not somatic breakpoints. Finally, a genotyping file was constructed with sequences of 12 bases adjacent to the breakpoint resembling the novel insertion and wild-type genome. If the wild-type sequence of a breakpoint was not known ("N"s in the reference genome used to align the bam file), the insertion was discarded.

**Step 3: Re-Genotyping**

Using pysam, reads spanning the region between both breakpoints of an insertion, extended by one base to either end, were extracted from the bam file and the sequence adjacent to each breakpoint identified. If more than 60 fragment were identified, the insertion was called as an artefact (high coverage). Otherwise, 12 bases of this sequence were compared to both the reference sequence and the sequence of the novel insertion by calculating an alignment score: scoring 1 point for matching bases, -2 points for mismatching bases and 0 points for any "N". In this way, each read was called as evidence supporting either the presence or absence of the insertion, depending on which alignment had the lower score. A score difference of 6 or less was required for a call. Reads having the sequence of both breakpoints simultaneously were marked as artefacts, as were reads matching neither reference nor novel insertion. An insertion is called "heterozygous" or "homozygous" if both breakpoints have at least one evidence read. "Homozygous" reads show no evidence of wild-type alleles. If there is only evidence for one of both breakpoints, the genotype is temporarily labelled "insertion?".

**Step 4: Combining genotypes**

Individual genotyping results were combined from all colonies from a single human donor or from all the related mice. Insertions with less than 10% wild-type colonies (most likely inherited through the germline), no "heterozygous" or "homozygous" calls, more than 10% artefactual calls, or more than 10% high- or no-coverage calls were discarded. The exact settings of this filter may be varied depending on the size of an experiment. If there are more "insertion?" than the sum of "heterozygous" and "homozygous" calls, the insertion is also discarded. Insertions are called potentially somatic if less than 50% of the colonies of an individual show evidence of the insertion, otherwise they were designated "germline", although for border-line cases these were manually checked for evidence of a very-high frequency somatic insertion (none were found in the mouse or human data).

**Step 5: Annotation**

The clipped parts of both breakpoints were aligned to the hs1 or GRCm39 genome separately and the coordinates are then lifted over to the assembly used to align the bam files. If the clipped part maps within 1000 bases of the breakpoint, the insertion was marked as "artefact". Additionally, the clipped part of each breakpoint is run against the DFAM database[3] using nhmmer[4] to identify the most likely family of retrotransposons  the insertion is most likely to belong to.  PolyA-Tails are identified by LBPs (left breakpoints) with a clipped part ending in at least 6 adenines or RBPs (right breakpoints) starting with at least 6 thymidines. An insertion is attributed to LINEs or SINEs if either both clipped parts map to the same family of LINE/SINE or if one end does and the other corresponds to a polyA tail. ERVs are called if both ends map to the same subclass of LTR.

#### Supplementary Method 2: Estimation of insertional mutation rates in mouse and human HSCs

Phylogenies were converted to ultrametric trees and the first 40 (mice) or 55 (human) mutations were clipped, since they occur *in utero*[5,6]. We then calculated the total number of HSC-years observed in a tree by summing remaining edge lengths (HSC time) scaled to the known age of the donor, assuming a constant rate of SBS after birth. The number of insertional mutations that occurred later than the above mentioned cutoffs ($n_{ins}$) were summed. Insertions with a timing that overlaps this cutoff were counted as the fraction of the edge after the cutoff divided by the total edge length and the sum of all fractions was rounded to the nearest number.

The following data was obtained for mice:

| Mouse | Age | $n_{ins}$ | Captured HSC time after birth (cell-years) |
|---|---|---|---|
| MD7180 | 30 months | 18 | 701 |
| MD7181 | 30 months | 7 | 230 |
| MD7182 | 30 months | 7 | 216 |

And for humans:

| Patient | Age | $n_{ins}$ | Captured HSC time after birth (cell-years) |
|---|---|---|---|
| AX001 | 63 years | 3 | 21,971 |
| SX001 | 48 years | 1 | 16,984 |
| KX008 | 76 years | 1 | 20,907 |
| KX004 | 78 years | 3 | 47,005 |
| KX003 | 81 years | 3 | 18,079 |
| KX002 | 38 years | 1 | 14,249 |
| KX001 | 29 years | 0 | 11,582 |

We then fitted a Poisson generalised linear model to estimate the rate of insertional mutations per HSC-year and obtained a rate of $2.79 \times 10^{-2}$ (95% confidence interval $1.93 \times 10^{-2} – 3.87 \times 10^{-2}$) insertional mutations per cell-year for mice and $7.96 \times 10^{-5}$ (95% confidence interval $4.26 \times 10^{-5}$ – $1.33 \times 10^{-4}$) for humans.

The number of stem cells at steady-state in humans is estimated to be 100,000 in humans[5] (95% confidence interval 20,000-200,000) and 70,000 in mice[6] (95% confidence interval 25,000-100,000). Assuming the number of stem cells has reached steady-state at birth for humans or 30-months for mice, by multiplying the rate of insertional mutagenesis per year and HSCs with the number of HSCs, a rate of insertional mutations per year in all stem cells of an individual was obtained, estimated to be 1838 per year (95% confidence interval 773-4334) for mice and 5.62 per year (95% confidence interval 1.54 - 17.72) for humans. By multiplication with the average expected natural lifespan (30 months for mice, 82 years for humans) the total number of insertional mutations expected during lifespan after birth was estimated to 4610 (95% confidence interval 1949-10,818) for mice and 460 (95% confidence interval 126-1456) for humans.

### Supplementary Method 3:

### Calculation of Recall and Precision for PEAR-TREE

The recall rate of PEAR-TREE (the number of true positives divided by the sum of true positives and false negatives) was estimated using heterozygous germ-line retrotransposon insertions detected in the donor tissue used in this study, but not present in the reference genome. The chance of discovering such a germ-line heterozygous insertion in each single colony indicates the chance of discovering a somatic single-colony insertion.

Insertional mutations where less than 10% of colonies could be identified as wild-type were considered germline. We determined the number of colonies of a mouse that the germ-line insertion had been discovered in during the initial discovery phase (steps 1 and 2 of Supplementary method 1). Based on the average of 71 heterozygous germ-line insertions in 10,424 individual discovery instances, heterozygous insertions had a chance of **54.8%** to be discovered in a single colony, which is expected to be equal to the recall for PEAR-TREE to detect a single-colony somatic insertion. We also analysed 23 homozygous insertions and found a recall of 84.5% for these. No differences were observed for recall between the three classes of retrotransposons.

Detection of the same insertional mutation in multiple colonies from the same individual was used to estimate the precision of PEAR-TREE (the number of true positives divided by the sum of true positives and false positives). Accurate detection of genuine insertional mutations is expected to result in the same mutation being shared by all colonies descended from the cell in which the mutation occurred, consistent with the phylogeny established using somatic single-base substitution mutations. False positive events are expected to appear in random colonies. For each multi-tip insertion, we calculated the probability that the pattern of colonies it was detected in could be explained by the null hypothesis, a random distribution (thus a false positive), $p_{random}$, as well as the probability that an insertion that is perfectly consistent with the phylogeny could occur by chance given the number of colonies carrying the insertion, $p_{min}$, using the following formulas

$$p_{min} = \frac{1}{\binom{N}{i}} \text{ and } p_{random} = \frac{\binom{C}{j}\binom{N-C}{i-j}}{\binom{N}{i}}$$

Where $\binom{n}{k}$ is the binomial coefficient $\frac{n!}{k!(n-k)!}$, N is the total number of colonies, i is the total number of colonies with the insertional mutation, C is the size of the best fitting clade and j is the number of insertional mutations inside the best fitting clade. For these calculations, colonies with evidence of the insertional mutation ("heterozygous", “homozygous” or "insertion?" genotypes), were considered as carriers of the insertion. Colonies where there was not sufficient evidence to be confident in a wild-type genotype were excluded.

The best clade was determined by finding the most recent common ancestor (MRCA) node of all "homozygous" or "heterozygous" colonies or its immediate ancestor, if this node would incorporate one or more "insertion?" colonies compared to the MRCA.

For some of the multi-colony insertions the number of colonies and/or the complexity of the associated phylogenetic tree were low enough that the pattern of insertions observed could reasonably be expected by chance ($p_{min} > 0.05$). These were be discarded from further analysis, because they cannot provide meaningful information about whether PEAR-TREE precisely detects true mutations in related colonies. Of the remaining multi-tip insertions, those with $p_{random} < 0.05$ were considered true positives, and the others true negatives.

Using this method on 104 considered multi-tip insertion with $p_{min} > 0.05$, 103 had a $p_{random} < 0.05$ and were thus considered true-positives, while one had an adjusted p-value of 0.11 and was thus considered a false-positive, resulting in a **precision of 99.0%.**

We verified the validity of this method to calculate the precision by re-applying the method after randomly shuffling the genotypes of every multi-tip insertion, leading to 3 of 104 mutations with $p_{min} < \alpha$, and a simulated precision under the null hypothesis of 2.9% which is less than $\alpha$.

### Supplementary Method 4: Testing the independence of insertional mutagenesis events in phylogenetic trees.

Several instances where multiple insertional mutations occurred together in the same branch of phylogenetic trees from humans and mice led us to hypothesise that insertional mutations do not occur independently from each other, but tend occur more frequently in related HSCs in the same subclades of the phylogeny. To test this, we simulated phylogenetic trees with insertional mutations distributed under the null hypothesis, where they are assumed to occur independently from each other.

Assuming that the structure of the phylogenetic tree based on SBS mutations is not disrupted by codependent somatic retrotranspositions, we used unaltered phylogenetic trees for this simulation. To simulate random, independent occurrence of insertional mutations across these phylogenies: for each insertional mutation, all branches that overlap the branch where the mutation occurred (in range of SBS mutations) were selected (See Figure S3a). The phylogenetic tree was then simulated many times, where this mutation was randomly assigned to one of the selected branches, weighted by the overlapping length divided by the total branch length. This method of randomisation retains the temporal distribution of insertional mutation across the donor's lifespan. For each donor, we simulated 50,000 phylogenetic trees by randomising all insertional mutations in this way (See example in Figure S3b).

To quantify correlated insertional mutations in each real and observed tree, where multiple mutations accumulate in the same colony, half of all insertions that occur in the same node as another mutation (double insertions) were summed and added to the number of insertions occurring in a direct child branch of a branch already harbouring an insertion (stacked insertions) (Fig S3 c).

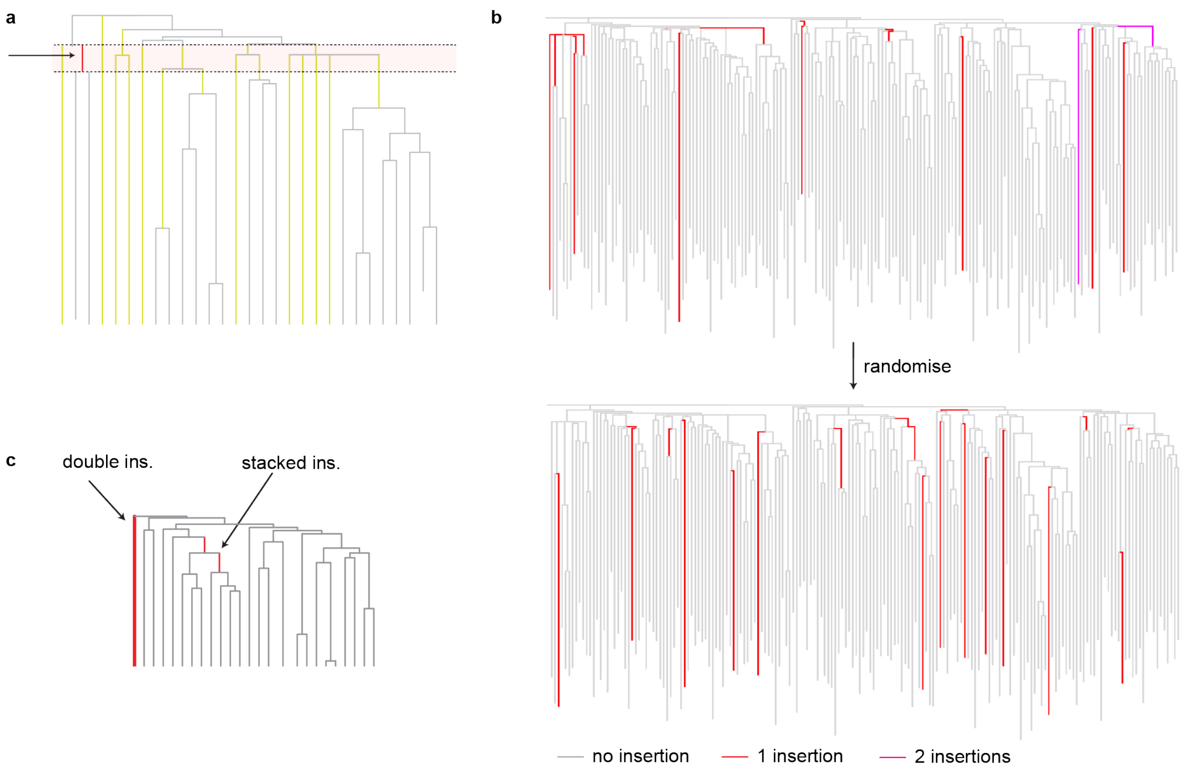


***Supplementary Figure S3****: (a) Method of selecting nodes to randomise insertional mutations. For each node with a detected insertional mutation (red, marked with an arrow), all nodes overlapping the SBS length were selected (yellow) and weighed by the length of each node overlapping the node with the insertion divided by the total length of the node. (b) An example of a randomised phylogeny generated for mouse MD7182. The observed tree is shown on the top, and one randomised tree generated from it on the bottom (c) Examples of double and stacked insertions, that were used to generate the test statistic.*

The p value was calculated as the quantile of the correlated mutation count determined from the observed phylogeny within the distribution of the 50,000 correlated mutation counts determined from the randomised trees.

The higher the number of insertions the higher the power to reject the null hypothesis. We only analysed phylogenetic trees with more than two insertions and obtained the following dataset:

| Species | Specimen | Age | Number of insertions | Correlated mutation count | Quantile (p-value) |
|---|---|---|---|---|---|
| Mouse | MD7180 | 30 months | 57 | 16 | 0.00008 |
| Mouse | MD7181 | 30 months | 10 | 4 | 0.0614 |
| Mouse | MD7182 | 30 months | 20 | 5 | 0.028 |
| Human | AX001 | 63 years | 3 | 1 | 0.015 |
| Human | KX003 | 81 years | 3 | 0 | 1 |

### Supplementary Method 5:

### PARTRIDGE: Detection of IAP-driven insertional mutagenesis in bulk bone marrow DNA

In contrast to LINEs and SINEs, retrotransposing ERVs insert into the genome with high fidelity, resulting in predictable sequences with two sequence-identical full-length long terminal repeats (LTRs) flanking the internal sequence. We developed a method to detect low-allele frequency ERV retrotransposition events, focussing on intracisternal A particle (IAP) ERVs, since expression of many IAPs were derepressed in HNRNPA1 2YF mice – **PARTRIDGE**: Precise Analysis of RetroTransposed Replications of IAPs by Deep Genomic Enrichment. Source code available on github (https://github.com/jeremydeuel/PARTRIDGE).

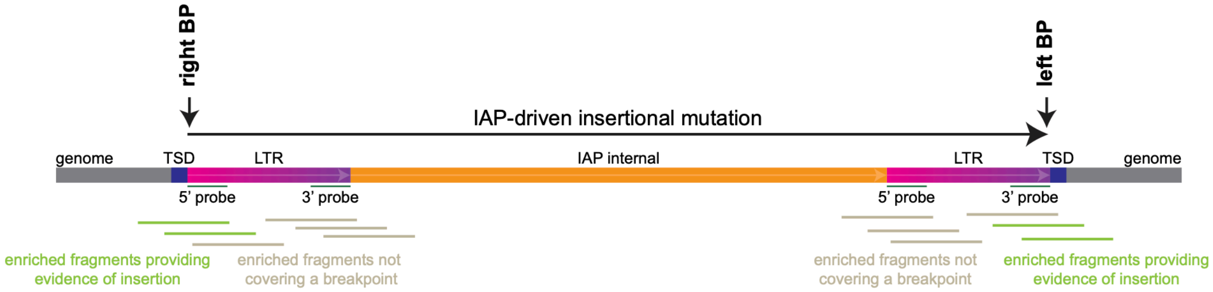


**Supplementary Figure S4**: Structure of an insertional mutation driven by an IAP ERV. ERVs insert with intact full-length long-terminal repeats (LTRs) on both sides (pink-purple), that are sequence-identical. The internal part (orange), encoding for viral proteins, can be full-length or truncated. We designed 120bp probes targeting the 3' and 5' end IAP LTRs, allowing enrichment of the LTR flanks, including chimeric fragments providing evidence for the novel insertion (lime green) spanning the breakpoints.

#### Hybridisation capture

Four biotinylated bait oligos were synthesised, targeting the 5' and 3' ends of the canonical LTRs annotated in DFAM (www.dfam.org) for "IAPLTR1_Mm", "IAPLTR1a_Mm","IAPLTR2_Mm" and IAPLTR2a":

Targeting 5' end of IAPLTR:

```
5'TGTTGGGAGCCGCGCCCACATTCGCCGTTACAAGATGGCGCTGACAGCTGTGTTCTAAGTGGTAAACAAATAATCTGCGCATGTGCCAAGGGTATCTTATGACTACTTGTGCTCTGCCT
5'TGTGGGGAGCCGCCCCCACATTCGCCGTTGCAAGATGGCGCTGACATCCTGTGTTCTAAGTGGTAAACAAATAATCTGCGCATGTGCCAAGGGTAGTTCTCCACCCCATGTGCTCTGCC
```

Targeting 3' end of IAPLTR:

```
5'TCTTGCTCTCTTGCTTCTTGCACTCTGGCTCCTGAAGATGTAAGTAATAAAGCTTTGCCGCAGAAGATTCTGGTCTGTGGTGTTCTTCCTGGCCGGTCGTGAGAACGCGTCTAATAACA
5'GCAATAGAGCTCTTGCTCTCTTGCTCTCTGGCTCCTGAAGATGTAAGCAATAAAGTTTTGCCGCAGAAGATTCCGGTTTGTTGCGTTCTTCCTGGCCGGTCGCGAGAACGCGTGTAAGA
```

Genomic DNA was extracted from bone marrow using Qiagen AllPrep Kits. Sequencing libraries were generated from 1 µg of DNA using the xGEN DNA Library Prep EZ UNI Kit with xGEN UDI-UMI full-length adapters according to the manufacturers protocol. Target fragment size was 350bp, and no PCR-Amplification was performed. Groups of sixteen libraries were consolidated into 2-4 µg pools of DNA. Hybridisation was performed with the xGEN Hybridisation Kit using xGen Blocking Primers for UDI-UMI full-length adapters and 4µl of pooled biotinylated bait probes at 1 µM final (total) concentration. Hybridisation was carried out at 65°C overnight (14h). After washing according to the manufacturer's protocol, captured sequences were amplified by 11 cycles of PCR and quantified with a fluorescent dye (Qubit). Libraries were sequenced on an Illumina Novaseq 6000 instrument using 250PE reads on an SE flow cell (two lanes).

**Building a HMMER-Model of IAP LTR edge sequences**

To accurately detect the ends of IAPs in sequencing data, we built a consensus model of the sequences at the edges of IAPs in the mouse genome that are most likely to retrotranspose. First, the genomic coordinates of 9228 IAP LTRs was determined, matching the Dfam models of IAP LTRs: DF000001785, DF000001786, DF000001787, DF000001788, DF000001789, DF000004145, DF000004162, and DF000004175. To identify pairs of nearby LTRs most likely to be active IAPs the following filters were applied:

- no further LTR between a pair of LTRs
- Size of IAP including both LTRs between 2-8 kb
- LTRs very similar (<5 edits per kb of sequence)
- TSD exactly 6bp
- LTR (5') starts with "TG" and ends (3') with "GA" or "CA".

Four HMMER (www.hmmer.org, version 3.4) models, representing quantitative consensus models of sequences at the ends of IAP LTRs, were built from the 3169 LTR pairs meeting these criteria. These contained the first and last 35bp of the 5' and 3' end, each in both forward and reverse complement orientations. Consensus sequences of these HMMER models were (generated with hmmemit, only forward sequences shown):

IAPLTR 5' end, forward

```
TGTAGGGAGCCGCGCTCACATTTGCCGTGGTATAAGT
```

IAPLTR 3' end, forward

```
CCTTCGCGCTTGGGGTCAAGACGAGTCTTTTAACA
```

### Detection of novel IAP insertions from hybridisation-capture sequencing data

Adapter sequences were trimmed from sequence data with Trimmomatic v 0.39 using standard settings. Reads were mapped to the mm39 reference genome using bwa-mem2 v 2.2.1 to generate bam files, sorted by coordinates. Reads where one end does not align to the reference genome were identified by filtering for soft-clipped ends longer than 24bp. The IAP LTR edge models were then compared to the entire read using nhmmer (through the python wrapper pyhmmer[7]). The best matching hmmer model was selected. Reads were discarded if they were clipped to the left and the model was 3' reverse or 5' forward or if they were clipped to the right and the model was 5' forward or 3' reverse, since these are not consistent with a bona fide IAP insertion (Figure S5a)

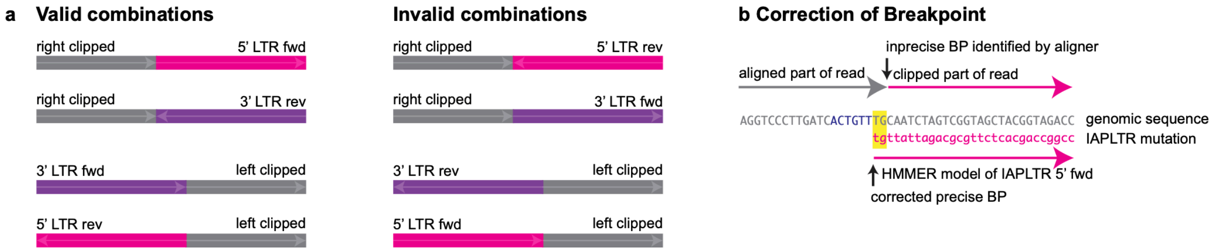


***Supplementary Figure S5:*** *(a) Only four of the eight possible combinations of clipping direction, LTR end and LTR direction (fwd or rev) are possible, the other four do not indicate a true insertion.*
*(b) Example of the correction of breakpoint: If the first couple of bases (in this example 2) of the clipped genomic reference sequence (gray) and IAPLTR (pink) overlap (yellow highlight), the aligner soft clips a read imprecisely relative to the start of an LTR. Using the HMMER model, this can be corrected by adjusting the breakpoint to the precise start of an IAPLTR.*

When the host genome sequence has – by chance – a small number of bases identical to the end of an IAP the breakpoint initially identified by the aligner did not precisely detect the start of the inserted IAP (Supplementary Figure S5b).  This breakpoint was therefore adjusted according to the start of the IAP LTR HMMER model allowing for up to 4bp correction.

After removing sequencing duplicates, breakpoints facing each other in opposite directions (consistent with an IAP insertion) were grouped within a 150bp sliding window and combined from all files. This list of breakpoint pairs was then used to re-genotype every BAM file. Insertion breakpoints found in more than one bone-marrow sample were labelled as germline and excluded from further analysis.

All raw reads covering the span between both breakpoints were then extracted from each BAM file, aligned and manually validated using the following criteria:

1. A least one supporting read with >20bp of LTR coverage for each end of the LTR is present
2. The insertion is not within an obviously repetitive region
3. No reads have discordant starting points of the LTR
4. There is a TSD of more than two bp and less than 30bp
5. Both LTR ends map to the same LTR in the reference genome (>93% sequence identity)- and this LTR is part of an LTR pair (2-8kb apart)

If the intersection of the results of both ends of an IAP insertion yields a single location in the reference genome, that insertion is marked as unambiguous source virus. One exception was made for an IAP present with two sequence-identical copies in C57BL/6 mice at chr3:60,397,294-60,402,505 and chr2:84,335,554-84,340,765. This virus caused the most insertional mutations in our experiments, and because the two loci are indistinguishable from each other, we treated chr3:60,397,294-60,402,505 as a unique locus. For other loci with ambiguous source loci, one of the possibilities was randomly selected as source locus to determine allelic frequency.

**Measuring allele frequency**

To estimate allele frequency we compared the number of reads associated with the somatic retrotransposition ($n_{destination}$) to the number of reads associated with the germline source ERV associated with the retrotransposition ($n_{source}$). Because these share the same LTR sequence, they will have been captured during hybridisation with identical affinity. Using samtools[8], $n_{source}$ was determined by counting uniquely mapping reads covering at least 25 bp of the source loci as well as 25bp of the adjacent genome. The same was done for the destination site $n_{destination}$, with the assumption that 25bp of coverage of the adjacent region would be enough to assert a mapping with mapq ≥ 40. The allelic frequency was then estimated as $AF = n_{destination} / (n_{source} + n_{destination})$. We validated these findings by measuring the AF of a germ-line novel insertion in the 1YF mouse line, and by simulation.

### Supplementary Method 6:

### Simulation of hybridisation capture enrichment

HMMER was used to align the four bait probes to the mm39 genome, identifying 48,290 potentially hybridizing loci. For each simulated experiment, 10 million reads were simulated using an R script. 80% of these were randomly picked to contain one of the loci identified by HMMER, weighted by the negative logarithm of the E-value (enriched fraction). 20% of the reads were generated by picking random locations from the entire mm39 genome (non-enriched fraction). To account for ligase artefacts, an additional 5% simulated reads were added, each with a random fragments from the enriched fraction concatenated to a random fragment from the non-enriched fraction. Novel insertions were simulated by adding one of ten IAP viral sequences identified as source of novel insertions into distinct positions of the genome and using the first and last 120bp of the LTR as enriched loci, simulating reads identically to the enriched fraction of the HMMER identified loci mentioned above. Read lengths were simulated to be approximately the same as in the observed samples by using a skewed normal distribution with location 250, scale 200, and shape 4, omitting results smaller than 51 bp length. Fragments shorter than 270 bp were padded to 270 bp length using xGen adapter sequences. The resulting sequences were piped into mason_frag_sequencing v 2.0.9[9] simulating 250PE Illumina reads. For each simulated experiment, 1000 novel ERV insertion were simulated with allele frequencies between 0 and 0.1. Additionally, 18 germ-line ERV insertions were simulated, that were the same for all experiments (9 heterozygous and 9 homozygous). Forty-three experiments were simulated and analysed with the same pipeline we used on the original samples. Precision and recall were calculated as previously described[10]
